# Distinct thermal responses of a host plant and an invertebrate herbivore affect ecosystem productivity and disease dynamics in a coastal marine ecosystem

**DOI:** 10.64898/2026.06.12.731926

**Authors:** Amy A. Briggs, Grace Callahan, Nicholas Yoong, John J. Stachowicz, Anya L. Brown

**Affiliations:** Dept. of Evolution & Ecology, University of California, Davis USA 95616; Center for Population Biology, University of California, Davis USA 95616; Northeastern University Marine Science Center, Nahant, MA 01908; University of California, Davis, Bodega Marine Lab, Bodega Bay, CA 94923

**Keywords:** climate change, disease ecology, interaction strength, microbes, seagrass, species interactions, thermal biology

## Abstract

Biological rates, like growth, tend to have unimodal (hump-shaped) responses to temperature, and these relationships can vary among species and biological processes. In most systems, full thermal performance relationships are rarely characterized for interacting species (e.g., consumer-resource or host-pathogen pairs), making it challenging to predict how their interactions, and subsequently, how communities, will shift with climate change. We investigated how the thermal responses of eelgrass (*Zostera marina,* an important marine foundation species in the N. hemisphere) and an isopod grazer (*Pentidotea resecata*), which putatively acts as an indirect vector of eelgrass wasting disease, interact to affect eelgrass productivity and wasting disease dynamics. In a laboratory experiment crossing five temperatures, two grazing, and two disease exposure treatments, across various metrics, eelgrass growth responded unimodally to temperature in the absence of grazers. Grazers depressed plant growth and flattened its thermal performance curves. Thermal performance curves for isopods indicated that increases in grazing and survival at intermediate temperatures negated concurrent gains in plant growth at these temperatures, while decreased isopod survival at high temperatures reduced their top-down effect on eelgrass. Isopods had negligible effects on plant disease responses, but warming reduced the time to disease onset and increased final disease severity. Overall, whole-plant disease severity remained low and did not substantially affect eelgrass leaf elongation, net growth, or rhizome dry mass. However, disease-treatment plants grew more new leaves at intermediate temperatures, possibly to combat losses in photosynthetic capacity in diseased leaf tissue. These results indicate that climate change-associated warming will likely increase eelgrass vulnerability to wasting disease. However, in sublethal outbreaks, disease could have less of an impact on eelgrass productivity than warming-induced increases in grazing. Thus, ignoring grazer responses to temperature could result in unreliable predictions of eelgrass productivity under climate change.

## Introduction

Infectious disease outbreaks can leave lasting legacies on ecological communities and affect many important ecosystem functions and services that humans derive from the environment. Climate change is predicted to promote disease outbreaks in many systems, often through direct and indirect temperature-mediated effects on hosts, pathogens, and disease vectors. For example, warming (which is projected to occur with climate change) can increase pathogen virulence (Kimes et al. 2012) and transmission (Elderd and Reilly 2014), affect vector competence–the capacity of a vector to transmit a disease (Paaijmans et al. 2012), and can make hosts more susceptible to infection (Gallana et al. 2013).

However, biological rates typically have a unimodal (hump-shaped) response to temperature (Dell et al. 2011), known as a thermal performance curve (TPC). Because of this, climate warming might have more complex effects on disease, potentially facilitating it under intermediate levels of warming, but having negligible effects or decreasing disease under higher levels (e.g., Mordecai et al. 2019, Shocket et al. 2020). Adding to this complexity, TPCs can vary among species (Deutsch et al. 2008), populations (King et al. 2018), and biological processes (Kingsolver et al. 2011, Sinclair et al. 2016), and can also be influenced by infection status (Greenspan et al. 2017, Padfield et al. 2020, Ware-Gilmore et al. 2021). The nonlinearity of TPCs and their variation among the processes (e.g., growth, immune response) and organisms (e.g., host, pathogens, and vectors) contributing to disease could therefore result in complex interactions under different environmental contexts.

However, for most infectious disease systems, particularly those that do not affect humans, we have limited information on how the main ecological actors (e.g., hosts, pathogens, vectors) that contribute to disease respond to temperature gradients, and how these unique thermal-response relationships interact to affect disease dynamics. Thus, unraveling these interactions is vital for forecasting the responses of ecosystems to climate change, including shifts in disease dynamics, and the cascading effects of these dynamics on other ecosystem outcomes, such as primary productivity or carbon storage.

Infectious disease outbreaks can have ecosystem-wide consequences when they affect habitat-building “foundation species”. For example, in the 1930s, a wasting disease induced a mass die-off of more than 90% of North Atlantic eelgrass, *Zostera marina* (Tutin 1942, Muehlstein 1989), resulting in dramatic shifts in nearshore ecosystems, declines in migratory waterfowl (Moffitt and Cottam 1941, Addy and Aylward 1944), and a collapse of several shellfish populations (Stauffer 1937) and local fisheries (Thayer et al. 1984, Short et al. 1987). Subsequent outbreaks of wasting disease have continued to threaten eelgrass across its broad geographic range, typically requiring multiple decades (or more) for recovery (Godet et al. 2008). Globally, seagrasses (including eelgrass) have experienced major declines, with disease as a major driver in many locations (Orth et al. 2006, De Los Santos et al. 2019).

As in many other disease systems, increased temperatures are hypothesized to play a role in eelgrass wasting disease outbreaks. Field studies have found that disease prevalence and severity increase in the summer (Groner et al. 2021) and during warm-temperature anomalies (Aoki et al. 2022, Graham et al. 2023). Additionally, the causative pathogen of eelgrass wasting disease, the slime-mold-like protist, *Labyrinthula zosterae*, grows more rapidly in culture at warmer temperatures (Dawkins et al. 2018). However, in laboratory infection experiments, warming does not significantly affect the likelihood of infection, and has inconsistent (and often negligible) effects on infection severity (Dawkins et al. 2018, Eisenlord et al. 2024). This inconsistency could arise because thermal biology would predict that warming could either increase, decrease, or have no effect on disease, depending on the magnitude of warming and the thermal performance curves of the organisms involved (e.g., hosts and pathogens). Additionally, pathogen performance measured in isolation does not necessarily predict within-host performance (Cohen et al. 2017), as the latter is a product of both the host and pathogen’s physiologies and their interaction.

Grazers could further complicate the response of this system to warming. Grazing by common eelgrass herbivores such as isopods, amphipods, and snails can indirectly promote eelgrass wasting disease, likely by creating wounds that facilitate pathogen invasion, suggesting that grazers might act as indirect disease vectors (Graham et al. 2024). Here, we use the term “vector” broadly, to encompass vectors that directly transmit pathogens, and those that indirectly facilitate disease through mechanical wounding or other actions. However, some grazer species, like the isopod *Pentidotea resecata*, will selectively consume diseased eelgrass tissue (Murray et al. 2024), whereas other taxa prefer healthy tissue (Graham et al. 2024). Therefore, grazers might facilitate or inhibit disease, depending on the grazer species that is present. How these processes interact under increasing temperatures, which could stimulate grazer metabolism, consumption, and movement (Lemoine et al. 2013, Cloyed et al. 2019), could further alter disease dynamics and influence the response of eelgrass systems to climate change.

Broadly, because of these (and other) effects of warming on ectothermic grazers, and potential asymmetries in the responses of grazers and primary production to warming, producer-consumer interactions are predicted to intensify under climate warming (O’Connor 2009, Hamann et al. 2021). As in host-pathogen systems, however, thermal sensitivities are rarely characterized for producer-consumer pairs, making it challenging to predict precisely how the sign and strength of their interactions will shift under different levels of warming. Additional complexity occurs when consumers act as direct or indirect disease vectors to their resource, which is common in many terrestrial and aquatic ecosystems, including many crop-pest systems (Eigenbrode et al. 2018, Nicolet et al. 2018, Gossner et al. 2021). Thus, a mechanistic understanding of how warming can alter these types of complex interaction networks will advance our ability to accurately forecast how such systems will respond to climate shifts.

Here, we use the eelgrass wasting disease system to assess how the thermal sensitivities of different host and grazer/vector traits interact to affect critical disease and ecosystem outcomes. We assessed these interactive drivers of disease through a laboratory mesocosm experiment assessing disease symptoms and plant growth in response to a fully factorial manipulation of temperature, disease exposure, and grazing. We also conducted separate grazing trials to measure grazing rates at different temperatures. This design enabled us to characterize the thermal performance of the eelgrass host and its herbivore/vector in isolation and in combination with each other, and to reveal emergent ecosystem effects on disease (infection onset time and disease severity) and eelgrass productivity (here, net growth rate), which mediates many of the ecosystem functions and services that eelgrass provides. We measured additional eelgrass responses such as final rhizome dry mass, aboveground vs. belowground biomass, and leaf production rate, to assess how warming, grazing, and disease exposure affect plant biomass and energy allocation to different tissue compartments. These metrics can reveal plant strategies for coping with stress and inform predictions of how climate change and disease will affect carbon cycling and storage in eelgrass ecosystems through changes in leaf turnover and litter production, as well as belowground biomass accumulation.

## Methods

### Field collections

We collected eelgrass and isopods at low tide in the eelgrass meadow near Westside Park in Bodega Bay (38°19.1920 N, 123°03.1890 W) on August 19-22, 2024. Diseased plants were kept separate from healthy plants until the experiment began. Disease was endemic in our system, and most plants in the field had lesions on older (third-rank or higher) leaves, as older leaves have had more time for disease exposure and lesion development. Therefore, for the healthy experimental plants, we removed all but the first two leaves and ensured no signs of disease were present (similar to Eisenlord et al. (2024)). For the “diseased” plants, we retained the first three leaves to acquire sufficient experimental plants with active lesioned tissue. Back at the lab, we inspected plants for disease again and then used a sterile blade to trim them to a consistent size. Rhizomes were trimmed to 4 cm. Leaves were trimmed to 15 cm, except for the third rank leaf of the non-focal, diseased plants, which was trimmed to 30 cm (which was necessary because most lesions occurred above 15 cm on the third rank). If leaves were less than 15 cm long, we trimmed the tip to ensure that every plant had a similar perimeter of damaged tissue through which a pathogen could potentially invade. After processing, plants were placed in a low-salinity (20 ppt), filtered (0.2 µm) seawater bath for two hours to reduce infection risk, as low salinity has been correlated with decreased infection severity (McKone and Tanner 2009, Brakel et al. 2019). Then they were placed in ambient salinity (34 ppt), filtered (0.2 μm) seawater at 13 °C to heal for one day.

### Acclimation

Following collection and processing, plants and isopods were separately acclimated to experimental temperature conditions over 4 days, to avoid acute temperature shock for individuals placed in the warmest treatments. Individuals were placed in tanks with flow-through water in the experimental water tables (water table setup below). Tanks were incrementally warmed or cooled as necessary until reaching their final treatment temperature on the 5^th^ day.

### Experimental mesocosm system

The mesocosm experiment crossed five temperature treatments (9.0, 15.8, 19.3, 22.6, 24.8 °C), two disease exposure treatments (exposure to a healthy vs. a diseased conspecific), and two grazing treatments (with and without isopod grazers). Each of these twenty treatment combinations had six replicate tanks (120 tanks in total), with each replicate containing one healthy focal plant that was used for all plant and disease response measurements, and either a healthy or a diseased conspecific (i.e., two plants per tank). Each tank (a 5-gallon, or 18.9 L bucket) received a constant supply of seawater, at an average rate of 40 L/hour (∼50 water turnovers per day) from a flow-through system with partial recirculation. Approximately 25% of the water was replaced with new water from the flow through seawater system at Bodega Marine Lab with each pass through the system. To minimize contamination by outside pathogens, the seawater underwent UV sterilization (36 W, Coralife Turbo-Twist 12x) immediately before entering the experimental tanks. Experimental temperature treatments were achieved with heaters or chillers associated with large water sumps that supplied water to each experimental tank and the surrounding water baths, which further maintained tank temperatures. Tanks in each temperature treatment were distributed between two water baths, and the entire experimental setup was distributed between two temperature-controlled rooms—a warm room maintained at 22 °C and a cold room maintained at 15 °C. Tank temperatures were monitored daily with a handheld temperature probe (Flinn Scientific), and with Hobo loggers—one logger was deployed in a random tank in each water table (i.e., two loggers for each temperature treatment), and recorded temperature every 30 minutes.

An airstone bubbler provided additional oxygenation and water movement in each tank. Experimental light levels were on average 15 μmol photons m^-2^ s^-1^ (±0.4 *SE*, *n* = 120; measured with a 2π PAR sensor on a diving PAM (Walz, GmbH) at the mid-depth in each tank), and were maintained using LED aquarium lights (Nicrew SkyLED Plus 2.0, Plant Light) on a 12:12 light:dark cycle.

### Mesocosm experiment

The mesocosm experiment lasted from August 26 to September 12, 2024. On the day that plants were placed in their experimental tanks, they were visually checked for disease, and given two needle punches through all leaves at the top of their sheath, to enable leaf elongation rates to be measured during the experiment (Westera and Lavery 2006). The wet mass of each plant was measured after spinning in a clean salad spinner for 15 s. Healthy plants were processed first to avoid cross-contamination, and sterile technique was used. After preparation, plants were planted in a pot with clean sand (Quickcrete playsand), to avoid contamination by pathogens present in their native sediment. Potted plants were then placed in their experimental tanks.

Isopods from each temperature acclimation group were measured (total length) and distributed among the tanks in the grazing and disease treatments in a stratified random manner to keep size structure similar among treatments. (Isopods were on average 19 mm ± 0.5 SEM.) At the end of the experiment, we collected isopods from each tank and counted to determine survival (after focal plants were removed). We initially placed four isopods per tank, but reduced density to two per tank on day four, due to unexpectedly high grazing damage in some treatments (leaving 120 isopods distributed across 60 tanks). In several tanks, an extra isopod was found at the end of the experiment (indicating there were a total of three isopods in the tank during the experiment). We assumed 100 percent isopod survival (three out of three), in these tanks.

### Disease measurements

To evaluate treatment effects on disease transmission and progression over time, we recorded each focal plant’s disease status (healthy/diseased) for each blade, every other day until the end of the experiment. Plant pots were attached by two monofilament lines to a square dowel rod placed atop the tank, out of the water. This created a hanging basket with a handle that the researcher could use to lift the plant for viewing without reaching into the water. Researchers also wore gloves and sterilized their hands in 70% ethanol between each plant during these surveys to minimize cross-contamination. After 15 days of exposure, we removed focal plants for final measurements and sample collections.

We then scanned the blades of each plant on an Epson Perfection V550 scanner at 600 dpi, and used an automated algorithm, the Eelgrass Lesion Image Segmentation Application, EeLISA (Rappazzo et al. 2021) to quantify the final diseased area (the sum of lesioned tissue area across all leaves) and disease severity (the proportion of total leaf tissue that was diseased) for each plant. We also recorded the presence or absence of dark lesions on rhizomes for each plant (ESM1 Figure S1). No plant rhizomes had lesions at the start of the experiment. Rhizome lesions had an unknown cause but could be necroses caused by drivers such as anoxia or indicate EWD infection, although, to our knowledge, no study has investigated lesion development in rhizomes due to wasting disease.

### Plant responses

Plant growth was measured as the change in whole-plant wet mass (a commonly used metric of net growth in studies on aquatic macrophytes) and the elongation of leaves (a metric of gross growth), standardized by the experimental duration. Wet mass was measured at the beginning and end of the experiment by spinning plants in a salad spinner for 15 s to remove excess water and then weighing the plant. Leaf elongation (areal growth of new leaf tissue) was determined from scanned photos of each plant’s blades collected at the end of the experiment, using ImageJ (Schneider et al. 2012) to measure the area (cm^2^) below the needle-punch scars on each leaf, summed across all the leaves on a plant, plus the area of all new leaves that grew during the experiment (e.g., (Westera and Lavery 2006, Graham et al. 2021). The rate at which new leaves were produced by a plant was calculated as the change in the total number of leaves associated with it between the beginning and end of the experiment, divided by the experimental duration. Seven plants (including two that died) had a leaf (or part of a leaf) detach during the experiment (spread across all disease exposure and grazing combinations; ESM Table S1), but this leaf material was included in the final leaf count and growth measurements.

To evaluate how different tissue compartments (e.g., belowground vs. aboveground biomass) might respond to the experimental treatments, we separated each plant into its constituent parts, e.g., rhizome, roots, sheath, new leaf growth (the area between the sheath and the needle punch holes, plus any new leaves produced since the start of the experiment), and old (original) leaf tissue (the leaf area above the needle punches). We dried samples at 60 °C, then weighed them to determine their final dry mass (g). Aboveground dry mass was calculated as the sum of the sheath, new leaf, and old leaf tissue. Belowground dry mass was the sum of the roots and rhizome tissue. Additionally, because rhizomes are a critical energy-storage organ in eelgrass (Burke et al. 1996, Touchette and Burkholder 2000), final rhizome dry mass was analyzed separately to infer how the treatments affected the plants’ energy reserves. To evaluate plant investment in belowground vs. aboveground growth at different temperatures, we also calculated the ratio of final belowground:aboveground dry mass.

### Eelgrass-grazer interaction strength

To estimate the top-down interaction strength (IS) of isopod grazers on eelgrass productivity, we calculated the natural log of the ratio of the mean final total dry mass of healthy treatment eelgrass in the absence 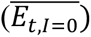 vs. presence 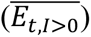 of isopods, divided by the experimental duration (Eqn. 1), based on equations of (Osenberg et al. 1997). Positive values indicate suppression of eelgrass biomass by grazers, with larger values indicating stronger top-down control.

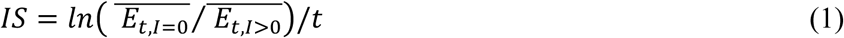

(We used final dry mass rather than the net growth rate to avoid undefined interaction strength estimates for plants that exhibited negative net growth, i.e., a loss of biomass.) We estimated the variance of each IS using the variance equation of (Osenberg et al. 1997), then calculated the 95% confidence intervals by multiplying the resulting standard errors by critical values from the t-distribution to account for small samples sizes.

### Grazing experiment

We measured how isopod grazing rates change with temperature using a separate, single-choice grazing experiment, conducted from September 3-13, 2024. Ninety-six isopods not used in the mesocosm experiment were randomly and evenly distributed among six temperature treatments ranging from 8.4 to 24.4 °C (ESM1 Table S3). After acclimation to their treatment temperatures (as in the mesocosm experiment), we placed individual isopods in separate experimental (140 mL) containers with mesh sides and lids that allowed water exchange. Containers were housed in the same temperature-controlled, flow-through water baths used in the mesocosm experiment, and grazing trials were conducted concurrently with the mesocosm experiment. At the start of each trial, isopods received a 2 cm^2^ section of an eelgrass leaf and were left to graze for five days. After this period, we collected and scanned the remaining leaf tissue, so its final area could be measured in ImageJ. Each temperature treatment also included 5-8 control (no-isopod) containers, to evaluate the change in leaf area in the absence of isopods. The isopod grazing rate was measured as the initial leaf area minus the final area, minus any scraped (i.e., not fully consumed) leaf area divided by two (Murray et al. 2024), standardized by the trial duration (cm^2^ d^-1^). Some isopods died or escaped, particularly in the warmest treatment, so we excluded those replicates from the final analyses, as we typically did not know when isopod grazing stopped. Each temperature had 11-16 replicates after excluding these isopods (ESM1 Table S4). Each isopod underwent two 5-day grazing trials, so to account for non-independence, we averaged their grazing rates across trials and fit thermal performance curves to these individual means.

### Analyses

#### Thermal performance responses

There is no universally best thermal performance curve model across traits, populations, or species (Kontopoulos et al. 2024), and the model used to fit data can strongly impact parameter estimates (Low-Décarie et al. 2017). Thus, we used the recommended approach of Kontopoulos et al. (2024) and fitted several TPC models that encompassed a range of shapes (including symmetric and asymmetric distributions) to each response variable. We then used AICc model comparison to select the best model. We chose models that had fewer parameters than temperature treatments (i.e., < 5) to reduce overfitting the data. We also selected models that could accommodate characteristics of the response variable, such as true negative values (e.g., net growth) or values constrained between zero and one (e.g., isopod survival, presence of rhizome necroses). Whenever possible for a response variable, we included models with biologically meaningful parameters, such as *r_max_* (the maximum rate) or *T_opt_* (the temperature at which *r_max_* occurs, i.e., the thermal optimum). For continuous response variables, such as eelgrass growth (change in wet mass, leaf elongation, final rhizome dry mass, and isopod grazing), we fit Lactin-2 (Lactin et al. 1995), Weibull (Angilletta 2006), and second-order polynomial (i.e., quadratic) models (ESM 1 Appendix 1), using the *rTPC* package (Padfield and O’Sullivan 2023) in R (R Core Team 2025). For binary responses (e.g., isopod survival and presence/absence of rhizome necroses), we fit binomial regression models with a logit link and second-order polynomial terms for temperature (Rebolledo et al. 2021, ESM1 Appendix 1), using base R functions. Rhizome necroses had many zeroes, especially in the lowest temperature treatment, so to minimize potential issues with separation when estimating parameter values, we used Firth’s bias reduction method (penalized likelihood; Heinze and Schemper 2002), computed with the *logistf* package (Heinze et al. 2025).

Eelgrass growth metrics in the grazing treatments did not exhibit typical unimodal thermal performance curves; instead, they increased monotonically with temperature. Models that could accommodate this relationship were fitted to these groups, including linear, logarithmic, power, and asymptotic regression models (equations in ESM1 Appendix 2). We used the same procedure as above to determine the best-fit model for each response.

Occasionally, for some plant growth responses, the best-fit model differed between disease treatments. When the second-best model in one treatment received similar support from the data as the best model (dAICc < 2), and matched the best model in the other treatment, we used this second-best model to fit the data in this group, which facilitated parameter comparison across disease treatments. This analytical choice did not qualitatively change the results for any response variables.

After determining the best-fit model for each response variable, we estimated key parameters describing the thermal curve for each experimental group, including the maximum performance (*r_max_*) for that response, its thermal optimum (*T_opt_*), and its thermal breadth (*T_br_*). *T_br_* was calculated as the temperature range at which a response is equal to or above 80% of its peak performance (Sinclair et al. 2016, Rebolledo et al. 2021). For models that did not include these parameters, we numerically estimated them from the fitted curves. *T_opt_*, *T_br_*, and *r_max_* were not calculated for responses that did not exhibit unimodal thermal performance curves (i.e., final disease severity or plant responses in the grazing treatments), as none of these responses were best fit by an asymptotic function or had an optimum.

To quantify uncertainty in non-linear model fits and around parameter estimates, we used bootstrapping (with 1000 bootstrap iterations) to estimate 95% confidence intervals (CI). For continuous variables, we used residual-resampling bootstraps; for binomial responses, we used case-resampling bootstraps to avoid boundary issues. Confidence intervals were calculated as the 2.5th and 97.5th percentiles of the bootstrap distribution.

#### Plant disease responses

Final (plant-level) diseased area and disease severity both increased with temperature. Based on the data and biologically feasibility, we fit linear, exponential, power, and second-order polynomial models to each disease exposure-grazing treatment group (equations in ESM1 Appendix 3). We then used model comparison to select the best-fit model and bootstrapping to quantify uncertainty, as described in the *Thermal performance* section. Prior to fitting the best model (exponential), we centered temperature on 15 °C, approximately the midpoint of our experimental temperature range and a typical spring-summer field temperature at our collection site. This centering facilitated ecological interpretation and stability of parameter estimates, with the parameter *a* representing the predicted response at this reference temperature.

The best-fit model was an exponential function, so we then used a generalized linear model (GLM) with log link function and a Gamma error distribution (to account for increasing variance with the mean) to test the effects of temperature, grazing, and disease treatments on diseased area. To accommodate zeroes, we added 0.04 to diseased area before fitting the GLM, which was ten percent of the smallest observed non-zero value and approximated our measurement precision. We compared models with different additive and interactive predictor structures using AICc, and estimated P-values for the best-supported model using the *car* package (Fox and Weisberg 2011). We checked model assumptions using the *DHARMa* package (Hartig 2026).

To test how treatments affected time to visible disease onset, we used the *survival* R package (Therneau 2024) to fit an accelerated failure time (AFT) model with a Weibull shape parameter to the disease incidence vs. day data, with grazing and disease treatments as categorical predictors, and temperature as a continuous predictor. We exponentiated the estimated coefficients for each predictor to calculate time ratios, which indicate the multiplicative effect of the predictor on time to disease onset (Therneau and Grambsch 2000). Thus, the AFT model allowed us to test how much the disease and grazing treatments altered time to disease onset relative to the reference treatment level for that predictor (exposure to a healthy plant, and no grazing, respectively), and how much a one-degree increase in temperature altered disease onset time. We evaluated model fit using its concordance index, which indicated how well predicted rankings of individual disease onset times matched observed rankings, with a value over 0.7 indicating an acceptable fit to the data. We compared models with different shape parameters (e.g., log-normal, log-logistic) and predictor structure, including additive and interactive models.

## Results

### Experiment treatments

Temperature treatments were relatively stable throughout the mesocosm and grazing experiments, with mean temperatures ranging from 8.4 to 24.8 °C (ESM1 Tables S2,S3, Figure S2). These temperatures spanned the annual range of monthly mean temperatures in Bodega Bay (∼10-18 °C; Reynolds et al. 2016, Sanford et al. 2019, Schiebelhut et al. 2023).

### Plant growth responses

Two plants in the hottest temperature treatment died during the experiment (one in the diseased-grazing, and one in the diseased-no grazing treatment). Their sheath and basal meristem rotted and leaves separated, so they were not included in analyses.

Plant growth responses, including the net growth rate (i.e., productivity, measured as change in wet mass d^-1^), leaf elongation rate (cm^2^ d^-1^), leaf production rate (leaves d^-1^); final aboveground, belowground, and rhizome dry mass (g); all showed qualitatively similar responses to grazing and disease exposure (Figures 1a-d, S4). Grazing had strong effects by altering the functional relationship between each of these responses and temperature. Specifically, no-grazing treatments exhibited classic, unimodal thermal performance curves (TPCs), with the best-fit model depending on the response (ESM2 Table S1). The thermal optimum for each no-grazing curve ranged between 15.8 and 20.8 °C, depending on the response and disease treatment (ESM1 Figure S4). (Parameter estimates are available in ESM1 Tables S5,S6.)

**FIGURE 1.**
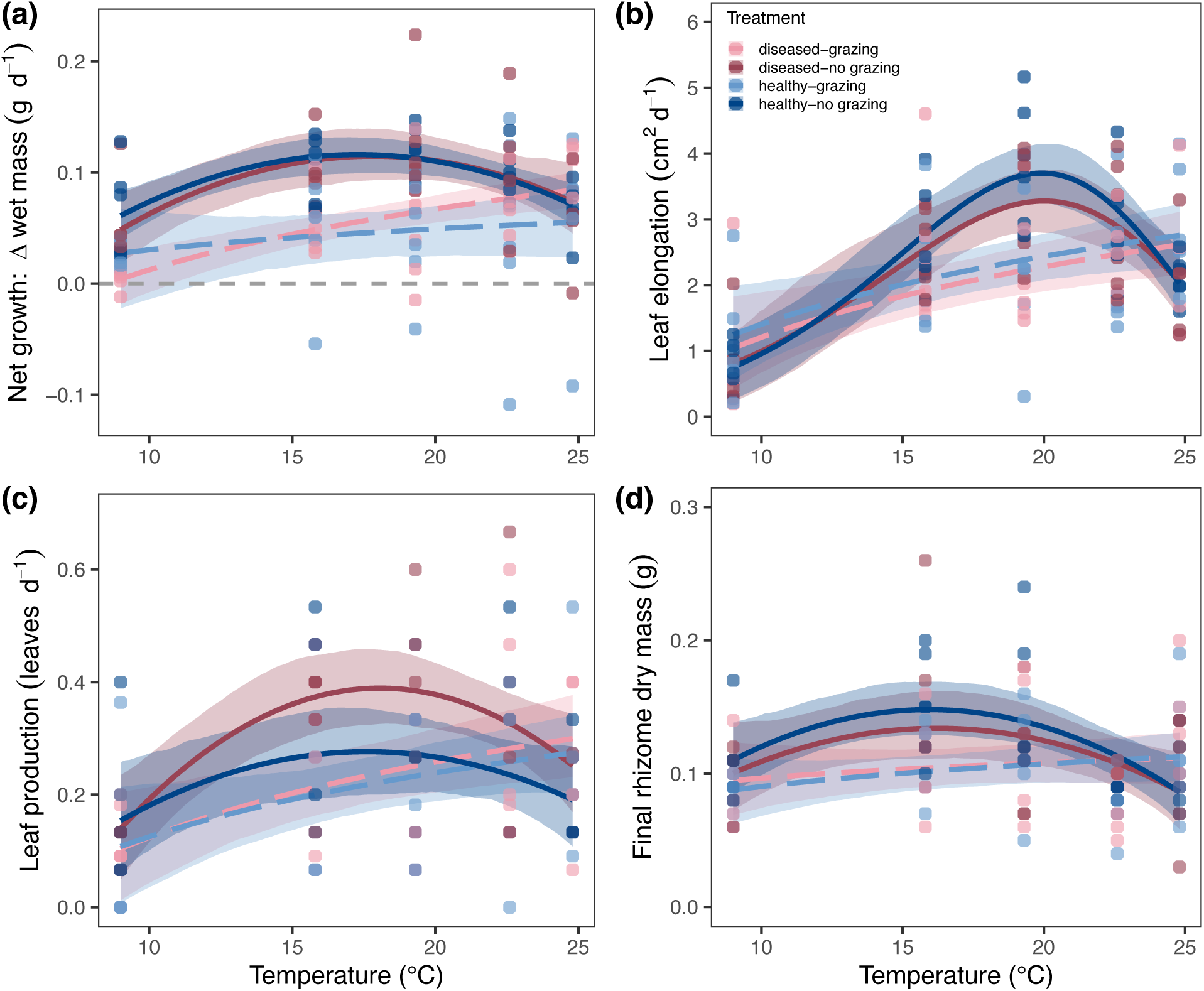
Plant thermal performance responses to grazing and disease exposure treatments in the mesocosm experiment. **(a)** Net growth rate (i.e., plant productivity): change in wet mass (g) per day for each plant. **(b)** Leaf elongation rate: area (cm^2^) of new leaf tissue produced per day by a plant **(c)** Leaf production rate (leaves d^-1^): rate at which the plant produced new leaves **(d)** Rhizome energy storage: final rhizome dry mass (g). Predicted model fits shown with 95 % CI. Points represent observed responses of individual plants. Color indicates treatment. Healthy treatment predictions are shown in dark and light blue, while diseased treatments are shown in red and pink. Grazing treatments are represented by the dashed colored lines.

In contrast, grazing treatments exhibited flattened, monotonically increasing relationships with temperature, best fit by a logarithmic function (ESM2 Table S2, ESM1 Table S7). Grazing depressed growth across all but the hottest (24.8 °C) and sometimes coldest (9.0 °C) temperatures, with the greatest reduction occurring near the thermal optimum (*T_opt_*) for the no-grazing treatments. For example, grazing reduced net growth at *T_opt_* from approximately 0.1 to ∼0.03 g d^-1^, a 70 % decrease. Thus, the top-down interaction strength between grazers and eelgrass, had a unimodal relationship with temperature (ESM1 Figure S6). Grazers had the most negative effects on eelgrass growth (i.e., the strongest top-down interaction strength) near its *T_opt_*, and weaker effects at both low and high temperatures.

Disease exposure (focal plants with a healthy vs. a diseased conspecific) generally produced smaller, more subtle effects on thermal responses than grazing. Across response variables, *T_opt_* was very similar across disease exposure treatments, with high confidence interval overlap (ESM1 Table S5). Thermal breadth, *T_br_*, and maximum performance (*r_max_*) also did not differ between disease treatments in the no-grazing groups for most responses (ESM1 Table S5). However, leaf elongation and leaf production rates both exhibited trends that could be ecologically meaningful. For leaf production rate, *r_max_* was higher in the diseased treatment (0.39 vs. 0.28 leaves d^-1^), a 39% increase relative to the healthy treatment. This difference is equivalent to growing ∼1.5 more leaves in two weeks at *T_opt_* in the diseased treatment vs. the healthy treatment, or ∼3 more leaves in a month. In contrast, maximum leaf elongation was higher in the healthy treatment (3.7 vs. 3.3 cm^2^ d^-1^), a 12% decrease with disease exposure.

### Rhizome necroses

In contrast to the plant growth responses, the likelihood that a plant had rhizome necroses at the end of the experiment had a unimodal relationship with temperature across all four grazing and disease treatment combinations (ESM1 Figure S7). The likelihood that a plant developed necroses peaked between 18.8 and 22.7 °C (depending on treatment) and declined in the hottest temperature treatment. Disease exposure did not substantially influence *T_opt_* or *p_max_*, the maximum probability of having rhizome necroses, but *T_opt_* estimates trended higher in grazing treatments (ESM1 Table S5).

### Disease responses

Disease transmission was high in the experiment—the majority of plants became diseased by the end of the experiment, including in the healthy treatment, indicating that pathogens were either introduced during the experiment, or present in or on the plants from the beginning (likely due to the very high prevalence of disease in the field during collections). However, disease severity (i.e., the proportion of lesioned tissue on an individual plant) was low, with lesions covering 16 percent or less of the total leaf area on plants (ESM1 Figure S8).

### Time to disease onset

An accelerated failure time (AFT) model with a Weibull shape parameter and without interactions was the best-fit model for the disease incidence vs. time data (ESM2 Tables S3,S4). This model had a concordance value of 0.72, indicating it fit the data reasonably well. Disease treatment and temperature significantly affected time to visible disease onset, but grazing did not have a significant effect (Table 1). Increased temperature accelerated disease onset, with disease occurring 2.5 % faster per 1°C increase in temperature (Table 1), so disease onset was ∼ 33 % faster in the hottest treatment relative to the coolest (which differed by 15.8 °C). This resulted in the median time to disease onset decreasing from of ∼8 days at 9.0 °C, to ∼6 days in the hottest, 24.8 °C treatment (Figure 2a). The diseased treatment (exposure to an actively diseased conspecific) also accelerated disease onset, decreasing the time to disease onset by ∼25%.

**FIGURE 2.**
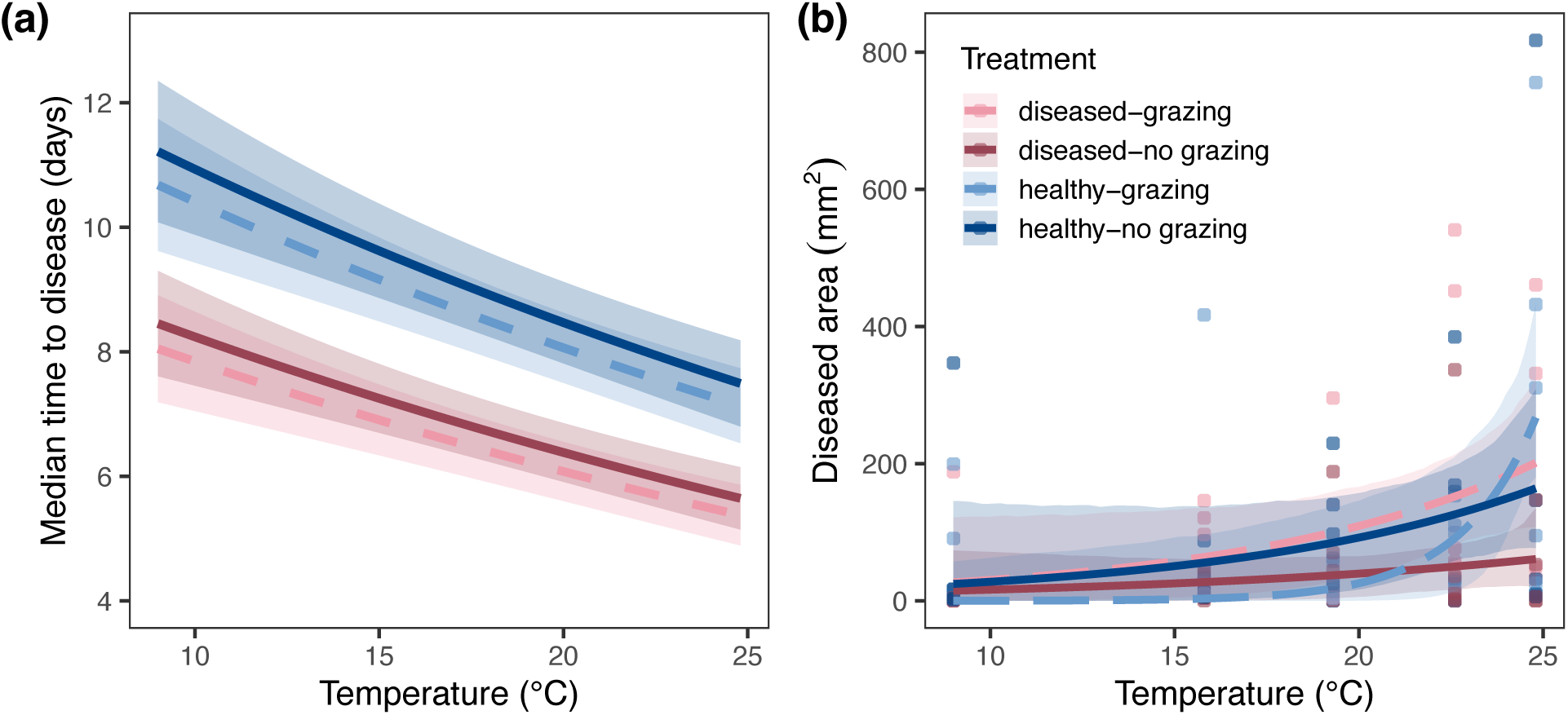
Disease responses to temperature in the mesocosm experiment **(a)** Predicted median time to disease onset (in days) vs. temperature for plants in the different disease exposure and grazing treatments, based on the AFT model results. See Table 1 for model coefficients. Shaded ribbons indicate SE for the predicted values. Dashed lines indicate grazing treatments; solid lines indicate no-grazing treatments. Healthy treatment predictions are shown in dark and light blue, while diseased treatments are shown in red and pink. **(b)** Final area (mm^2^) of diseased tissue on plants in the mesocosm experiment. Lines show best-fit model predictions, with 95% CI. Points indicate responses of individual focal plant replicates.

**TABLE 1.** Results for the best-fit accelerated failure time model of disease onset vs. time, which included main effects of disease exposure, grazing, and temperature (but no interactions). The time ratio (TR) indicates the proportional change in time to disease onset relative to the reference treatment level for categorical variables. (Reference levels were healthy for the disease treatment, and no-grazing for the grazing treatment.) The TR for temperature, a continuous covariate, indicates the relative change in time to infection for a 1-degree increase in temperature. TRs < 1 indicate a decrease in time to disease onset.

| Predictor | $\beta$<br>(log-time) | Time Ratio | 95% CI | P-value |
| --- | --- | --- | --- | --- |
| **Intercept** | **2.802** | **16.477** | **[12.152, 22.34]** | **<0.001** |
| **Treatment: disease** | **-0.283** | **0.754** | **[0.643, 0.883]** | **<0.001** |
| Treatment: grazing | -0.049 | 0.952 | [0.813, 1.115] | 0.542 |
| **Temperature (°C)** | **-0.026** | **0.975** | **[0.96, 0.989]** | **<0.001** |

### Final disease severity

Both final area of lesioned tissue (summed across all leaves on a plant) and final disease severity (proportion of total leaf area lesioned) increased exponentially with temperature (Figures 2b, S8; ESM2 Table S5). Confidence intervals for the predicted fits and parameter estimates substantially overlapped across the four disease exposure and grazing treatment groups, suggesting weak or no evidence for differences among groups (ESM1 Table S8). GLM results supported this conclusion—temperature significantly increased diseased area (p < 0.001), while disease and grazing treatments and their interaction had no significant effects (ESM1 Table S9). This pattern was consistent across all candidate model structures. The exponential rate coefficient, *b*, describing the proportional change in lesion area per °C, indicated that lesioned area doubled with every 1.6-5.3 °C increase in temperature (ESM2 Table S8), depending on the grazing and disease group (a 13-53 percent increase per °C). For the pooled data (across all treatment groups), *b* = 0.145 [95% CI: 0.063, 0.263], corresponding to a 16 percent increase per 1 °C increase, or a doubling every 4.7 °C (ESM1 Figure S9).

### Isopod responses

Like eelgrass, isopods exhibited unimodal responses to temperature (Figure 3). Isopod survival had a *T_opt_* of 13.8 and 15.4 °C (in diseased vs. healthy plant disease exposure treatments, respectively), after which it sharply declined, with one hundred percent mortality in the hottest 24.8 °C treatment. However, plant disease exposure treatments did not strongly affect isopod survival TPCs (ESM1 Tables S11, S12). Isopod grazing, measured in the separate grazing experiment, was best-fit by a second-order polynomial model (ESM2 Table S7), and had a *T_opt_* of 15.4 °C (ESM 1 Table S10).

**FIGURE 3.**
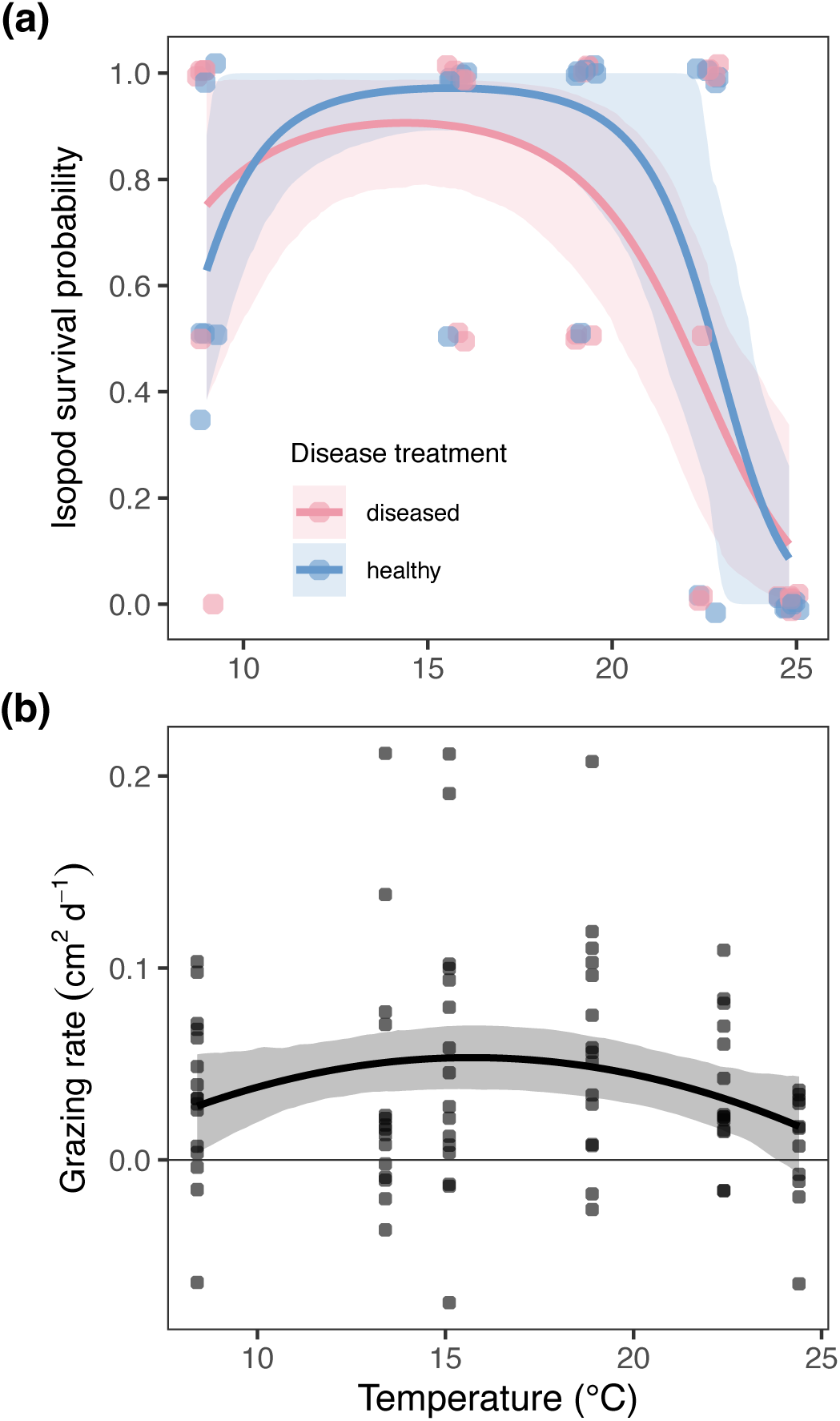
Isopod thermal responses **(a)** Isopod survival in different plant disease exposure treatments (colors) in the mesocosm experiment, and **(b)** isopod grazing rates measured in a separate grazing experiment. Panels show the best-fit lines with 95% CI. Points indicate observed responses of individual replicates (the proportion of surviving isopods in each tank for **a** or the mean grazing rate measured across two trials for each isopod in **b**). A small jitter was added to points in Panel **a** so that overlapping points could be seen.

## Discussion

Here, through manipulative experiments we demonstrated that temperature has critical, non-linear effects on eelgrass growth and morphological metrics (e.g., rhizome dry mass, below:aboveground ratio), grazing rates and grazer survival, as well as eelgrass wasting disease onset time and severity. Eelgrass and isopods varied in their sensitivity to temperature. This resulted in complex interactive effects of temperature and grazing on net primary productivity (i.e., eelgrass net growth). Eelgrass responses also varied in their reaction to disease exposure. However, disease had small effects on most eelgrass responses (e.g., net growth or rhizome dry mass), especially compared to the effects of warming or grazing, indicating that plant growth demonstrated some tolerance to wasting disease, at least in the short term. Overall, the different responses of eelgrass metrics to warming temperatures and/or disease suggest compensatory strategies that plants employ to tolerate these stressors and mitigate their impacts on host performance. We discuss these strategies and their potential limitations in greater detail below.

### Temperature effects on plant-herbivore interaction strength and eelgrass productivity

Isopod responses (grazing and survival) peaked at similar, but slightly lower temperatures (∼1-2 °C) than most eelgrass responses. As a result, heightened grazing pressure at intermediate temperatures cancelled out most gains in eelgrass growth, flattening the net growth curve of eelgrass in the grazing treatment (Figure 1, ESM S4). Although we observed low per capita grazing rates during the grazing trials (potentially due to leaf section size and/or differences in isopod risk perception between the experiments (Teckentrup et al. 2018, Murray et al. 2024) the general shape of the grazing curve qualitatively matched isopod survival in the mesocosm experiment, and our expectations based on the eelgrass net growth curves.

Because respiration is generally more sensitive to warming than photosynthesis (López-Urrutia et al. 2006, Yvon-Durocher et al. 2010), climate change-associated warming is predicted to strengthen herbivore-primary producer interactions, making plant biomass increasingly limited by herbivory (O’Connor 2009, Hamann et al. 2021). Here, our experiments illustrate how distinct thermal sensitivities of a primary producer (eelgrass, *Z. marina*) and its herbivore (isopod, *P. resecata*) give rise to a unimodal relationship between top-down interaction strength and temperature, such that grazers increasingly suppress plant growth with moderate temperature increases, but have decreasing top-down control with larger temperature increases. As a result, grazers caused net primary productivity and the standing stock biomass of eelgrass to increase slowly and monotonically with warming, in a manner that would not be predicted by the response of eelgrass to warming in the absence of grazers.

The unimodal relationship between the top-down interaction strength and temperature was driven, in part, by the lower thermal tolerance of isopods to high temperatures compared to eelgrass. Isopods had declining grazing rates in the 22.6 °C treatment and could not survive at 24.8 °C (Figure 3), while eelgrass maintained positive, albeit declining, growth for most response metrics at the same temperatures (Figure 1). Ectothermic herbivores have lower upper thermal limits than their resource in a variety of systems, e.g., intertidal gastropods and algae (Mertens et al. 2015), and aquatic zooplankton and phytoplankton (West and Post 2016). Such a decline in thermal tolerance with trophic level can occur through a variety of mechanisms, including the differential thermal sensitivities of respiration and photosynthesis discussed above, as well as faster increases in consumer metabolism with temperature than ingestion rates (Álvarez-Codesal et al. 2023), and declines in the palatability and/or nutritional quality (e.g., C:N) of producers with warming (Bauerfeind and Fischer 2013, Hernán et al. 2017, Wan et al. 2023). Although we do not know the mechanism here, we can infer that eelgrass net primary productivity (and hence standing stock biomass) would be released from top-down limitation under sufficient climate warming (assuming other consumers with a higher thermal tolerance are not present). This vulnerability of isopods to high temperatures could enhance the resilience of eelgrass to heat stress (e.g., during the summer or warm-water events) by reducing negative grazing impacts.

### Variation in thermal responses of different eelgrass traits

Despite the qualitatively similar responses of different eelgrass growth metrics to warming, there were some distinctions that suggest mechanisms that plants use to tolerate thermal stress. Our estimated thermal optima for different eelgrass growth responses ranged between 15.8-19.8 °C, which falls within the range observed for other northern eelgrass populations (Lee et al. 2005, Nejrup and Pedersen 2008, Niu et al. 2012, Beca-Carretero et al. 2018). However, rhizome dry mass, a metric of carbon storage and plant energy reserves, had a lower *T_opt_* than other plant responses like leaf elongation and net growth (Figure S4a). Previous work has found that eelgrass allocate less energy to rhizome storage and more energy to maintenance and growth under heat stress (Moreno-Marín et al. 2018). This might occur because, like in many other primary producers, eelgrass respiration tends to increase faster with warming than photosynthesis (Marsh et al. 1986). This discrepancy could shrink the energetic budget of eelgrass, thereby reducing the availability of carbon for storage in tissues like rhizomes. Our results illustrate that this process could begin at temperatures below the threshold at which negative effects of warming can be seen in more commonly measured metrics of eelgrass growth, such as change in total biomass or leaf elongation. Furthermore, the greater sensitivity of rhizome biomass to warming compared to other plant responses suggests that reducing energy allocation to rhizome storage or drawing on energy reserves from rhizomes might enable plants to maintain other forms of growth. This response could be beneficial in the short-term, allowing the plant to maintain photosynthetic capacity by continuing leaf growth. However, compensatory responses that allow an organism to tolerate stress can come at a cost to future performance and fitness (Sokolova 2021), sometimes even after the removal of the stressor (Martínez-De León and Thakur 2024). Thus, reduced energetic reserves in eelgrass rhizomes could limit the plants’ future reproductive output and growth (Govers et al. 2015) or make plants more sensitive to additional stressors, such as low light levels in deeper or turbid habitats (Lefcheck et al. 2017, Aoki et al. 2020), high herbivory pressure, or during extended or recurring heat waves (Nejrup and Pedersen 2008). Indeed, negative effects of warming on eelgrass growth can manifest weeks to months after heating occurs (Reynolds et al. 2016).

### Effects of temperature on eelgrass wasting disease

Observational field studies have documented correlations between eelgrass wasting disease prevalence and severity with warmer temperatures (Groner et al. 2021, Aoki et al. 2022, Graham et al. 2023). Here, we provide the first robust experimental evidence for the shape of this relationship—diseased tissue on individual plants increases exponentially with temperature, while time to disease onset decreases proportionally (Figure 2). Thus, warming has multiplicative (rather than additive) effects on disease. These results have several important implications. First, they suggest that the pathogen (*L. zosterae*) performs relatively well at high temperatures, especially compared to the host. Second, eelgrass wasting disease could spread more quickly under warmer temperatures, accelerating the emergence of outbreaks. Therefore, we might expect eelgrass populations to experience greater cumulative disease severity over time, as infections have more time to spread and progress within individual plants while conditions are favorable. (The exponential increase in diseased tissue in plants at the end of the experiment (Figure 2b) supports this inference at the individual level.) Additionally, these multiplicative effects could facilitate abrupt shifts in disease dynamics during extreme heat events, potentially pushing populations past ecological tipping points.

### Effects of disease on eelgrass productivity

Despite the effect of warming on disease onset time and lesion size, our experiment showed that across temperatures, disease exposure had a relatively small effect on eelgrass productivity, especially compared to other factors like grazing. In contrast, Graham et al. (2021), found that individual eelgrass blades grew more slowly when they were diseased, and diseased plants had reduced sugar storage in rhizomes. The lack of substantial differences between disease exposure treatments could reflect the low disease severity of most infected focal plants in our study, and the fact that some plants in both treatments developed signs of disease, albeit at different rates.

Additionally, the eelgrass exhibited other nuanced responses to disease exposure, indicating that plants altered allocation of resources to minimize disease impacts at the whole organism scale. For example, plants produced new leaves faster (especially at the *T_opt_* for growth) when exposed to diseased conspecifics (Figure 1c), while they generally maintained shoot growth (leaf elongation; Figure 1b). This suggests that plants might begin to invest more in the growth of new, uninfected leaf tissue while reducing investment in diseased leaf tissue, to maintain overall photosynthetic capacity, since disease reduces photosynthetic capacity of leaves (Ralph and Short 2002). Reallocation of resources between uninfected and infected tissues is a widely-documented compensatory strategy that plants use to deal with infection (Pagán and García-Arenal 2020). Thus, together with the results of Graham et al. (2021), our results illustrate how the unit of measurement (e.g., growth of a single leaf vs. whole-plant growth) can reveal different aspects of plant responses to infection.

We observed similarly low plant-level disease severity in our experiment as in other studies (Jakobsson-Thor et al. 2018, Schenck et al. 2023, Agnew-Camiener et al. 2025). However, more substantial negative effects of disease (on growth or reproductive output) might require longer timescales to manifest than our experiment could capture, as plants deplete energetic reserves and other resources to combat infection. *Labyrinthula* spp. (pathogen) virulence also varies among plant populations and pathogen strains (Trevathan-Tackett et al. 2018, Dawkins et al. 2018, Agnew-Camiener et al. 2025), with unknown effects on eelgrass performance and fitness.

### Grazers effects on plant-level wasting disease

Beyond temperature, grazing was another factor that we hypothesized would influence disease dynamics in our system. However, despite the strong effects of grazing on eelgrass growth, the grazer *P. resecata* did not significantly affect disease onset time or final lesioned area (Figure 2a,b). Eelgrass wasting disease is primarily transmitted via water, and relatively low densities (e.g., ∼6 cells/mL) of the pathogen *L. zosterae* can induce infection (Eisenlord et al. 2024). Therefore, high rates of environmental transmission might outweigh any effects of grazers on disease transmission, and subsequently, onset time.

In contrast to our results, field studies by Aoki et al. (2025) and Graham et al. (2024) found positive correlations between grazing scars and eelgrass wasting disease, which authors suggest indicates that grazing promotes disease. However, given that *P. resecata* preferentially grazes on diseased over healthy eelgrass tissue (Murray et al. 2024), the positive association between grazing scars and disease observed in the field could reflect grazer attraction to diseased leaves, rather than grazing promoting infection. At the plant-level, the net effect of grazing on disease severity might depend on the total grazing pressure (a product of grazer density and per capita grazing rates), and grazer identity, since species can vary in their preference for diseased tissue, and in the type of damage they inflict (Graham et al. 2024). Therefore, the grazer-disease relationship in eelgrass is likely context-dependent, and grazing could be a less important driver of wasting disease dynamics than previously suggested.

Overall, our results indicate that increased disease burdens on eelgrass under future climate warming might have relatively small effects on most eelgrass growth responses compared to the effects of increased grazing. Therefore, depending on grazer densities and other moderating factors, eelgrass standing stock biomass and belowground biomass at our sites might remain similar to current conditions, particularly under modest climate warming scenarios (e.g., a 2-degree increase in average summer temperatures). However, because warming-enhanced grazing could increase flux of eelgrass carbon into foodwebs, ecosystem functions like carbon cycling could shift substantially in eelgrass beds with climate change, even when standing stock biomass appears stable.

### Predicting outcomes of species interactions under climate change

Together, our results underscore how considering nonlinear effects of temperature on species interactions is likely critical to predicting community and ecosystem responses to a changing climate, as effects of warming on one organism (like a plant host) cannot be understood in isolation from their interactions with others (like grazers and pathogens). The shape characteristics of the thermal performance curves (TPCs) of interacting species will determine the thermal response of their interaction strength. For example, when herbivores exhibit similar or lower tolerance to high temperatures relative to their resource (e.g., when the herbivore TPC has a similar or lower thermal maximum than their resource), this can produce a unimodal relationship between top-down interaction strength and temperature (ESM 1 Figure S10). In contrast, when herbivores are much more tolerant to warm temperatures (e.g., when herbivores have a *T_opt_* that is higher than the thermal maximum of their resource), this could lead to monotonically increasing interaction strength with temperature (Figure S10). Where current and future environmental temperatures fall along these organisms’ TPCs will therefore affect how interaction strength will shift in different systems (either increase, decrease, or stay the same).

Ectothermic consumers are already close to their thermal optima and upper thermal limits under current conditions in a variety of systems (Pincebourde and Casas 2019), especially at lower latitudes (Deutsch et al. 2008). Additionally, upper thermal limits tend to decline with trophic position (Da Silva et al. 2023). Thus, a relatively small degree of climate warming could reduce the top-down interaction strength between herbivores and producers and even release net primary productivity from top-down control, potentially allowing producer standing stocks to increase in these systems. In contrast, higher latitude (temperate) consumers generally experience temperatures well below their thermal optimum (Deutsch et al. 2008), indicating that climate warming could intensify interaction strength and their top-down control of net primary productivity, thereby preventing producer biomass from increasing. These shifts could therefore alter the biogeographic pattern of increasing herbivore-producer interaction strength with decreasing latitude (Zvereva and Kozlov 2021). Of course, variability in thermal responses among other community members (competitors, predators, mutualists, etc.), as well as indirect effects of warming (e.g., shifts in herbivore body size), could modify these general patterns, but they nonetheless provide a useful baseline for predicting how herbivore-producer interaction strength will shift with rising temperatures.

A similar TPC-based framework can also apply to host-pathogen interactions. Mismatches in host and parasite thermal performance (e.g., a parasite with a lower *T_opt_* than its host) can drive parasite extinction under modest warming, effectively releasing hosts from harmful parasite effects (Gehman et al. 2018). This example is analogous to the release of eelgrass growth from grazing pressure that we observed at high temperatures. Together, these examples illustrate that the magnitude and direction of warming effects on biotic interaction strength, whether herbivory or parasitism, are contingent on the thermal performance curves of the organisms involved, and that the same framework can apply across different types of ecological interactions. Since species interactions are critical for shaping community composition and stability (McCann et al. 1998), applying this approach to different systems and across larger networks of species interactions will improve our ability to forecast how climate change will reshape communities and the ecosystem functions that they support.

## Supporting information

Supplemental model comparison tables

## Acknowledgements

We thank A. Lawson, J. Wilson, A. Leal, C. Hundley, M. Rajagopalan, C. Sorensen, and N. Yau for their help with field and/or lab work, as well as Bodega Marine Lab, especially the Aquatic Resource Group and Physical Plant staff. We thank the Center for Population Biology Postdoctoral Fellowship at UCD (to AAB) and NSF grants 23-11578 (to JJS), UC Davis start-up funds (to ALB) for funding that supported this study.

## References

1. Addy, C. E., and D. A. Aylward. 1944. Status of Eelgrass in Massachusetts during 1943. The Journal of Wildlife Management 8:269.

2. Agnew-Camiener, M. V., M. E. Eisenlord, C. S. Friedman, H. J. Schreier, and C. A. Burge. 2025. Pathogenicity and phylogeny of *Labyrinthula* spp. isolated in Washington and Oregon, USA. Journal of Eukaryotic Microbiology 72:e13073.

3. Álvarez-Codesal, S., C. A. Faillace, A. Garreau, E. Bestion, A. D. Synodinos, and J. M. Montoya. 2023. Thermal mismatches explain consumer–resource dynamics in response to environmental warming. Ecology and Evolution 13:e10179.

4. Angilletta, M. J. 2006. Estimating and comparing thermal performance curves. Journal of Thermal Biology 31:541–545.

5. Aoki, L. R., K. J. McGlathery, P. L. Wiberg, and A. Al-Haj. 2020. Depth Affects Seagrass Restoration Success and Resilience to Marine Heat Wave Disturbance. Estuaries and Coasts 43:316–328.

6. Aoki, L. R., B. Rappazzo, D. S. Beatty, L. K. Domke, G. L. Eckert, M. E. Eisenlord, O. J. Graham, L. Harper, T. L. Hawthorne, M. Hessing-Lewis, K. A. Hovel, Z. L. Monteith, R. S. Mueller, A. M. Olson, C. Prentice, J. J. Stachowicz, F. Tomas, B. Yang, J. E. Duffy, C. Gomes, and C. D. Harvell. 2022. Disease surveillance by artificial intelligence links eelgrass wasting disease to ocean warming across latitudes. Limnology and Oceanography 67:1577–1589.

7. Aoki, L. R., C. J. Ritter, D. S. Beatty, L. K. Domke, G. L. Eckert, O. J. Graham, C. P. Gomes, C. Gross, T. L. Hawthorne, E. Heery, M. Hessing-Lewis, K. Hovel, K. Koehler, Z. L. Monteith, R. S. Mueller, A. M. Olson, C. Prentice, B. Rappazzo, J. J. Stachowicz, F. Tomas, B. Yang, C. D. Harvell, and J. E. Duffy. 2025. Seagrass wasting disease prevalence and lesion area increase with invertebrate grazing across the northeastern Pacific. Ecology 106:e4532.

8. Bauerfeind, S. S., and K. Fischer. 2013. Increased temperature reduces herbivore host-plant quality. Global Change Biology 19:3272–3282.

9. Beca-Carretero, P., B. Olesen, N. Marbà, and D. Krause-Jensen. 2018. Response to experimental warming in northern eelgrass populations: comparison across a range of temperature adaptations. Marine Ecology Progress Series 589:59–72.

10. Brakel, J., S. Jakobsson-Thor, A.-C. Bockelmann, and T. B. H. Reusch. 2019. Modulation of the Eelgrass – Labyrinthula zosterae Interaction Under Predicted Ocean Warming, Salinity Change and Light Limitation. Frontiers in Marine Science 6:268.

11. Burke, M. K., W. C. Dennison, and K. A. Moore. 1996. Non-structural carbohydrate reserves of eelgrass Zostera marind. Marine Ecology Progress Series 137:195–201.

12. Cloyed, C. S., A. I. Dell, T. Hayes, R. L. Kordas, and E. J. O’Gorman. 2019. Long-term exposure to higher temperature increases the thermal sensitivity of grazer metabolism and movement. Journal of Animal Ecology 88:833–844.

13. Cohen, J. M., M. D. Venesky, E. L. Sauer, D. J. Civitello, T. A. McMahon, E. A. Roznik, and J. R. Rohr. 2017. The thermal mismatch hypothesis explains host susceptibility to an emerging infectious disease. Ecology Letters 20:184–193.

14. Da Silva, C. R. B., J. E. Beaman, J. P. Youngblood, V. Kellermann, and S. E. Diamond. 2023. Vulnerability to climate change increases with trophic level in terrestrial organisms. Science of The Total Environment 865:161049.

15. Dawkins, P., M. Eisenlord, R. Yoshioka, E. Fiorenza, S. Fruchter, F. Giammona, M. Winningham, and C. Harvell. 2018. Environment, dosage, and pathogen isolate moderate virulence in eelgrass wasting disease. Diseases of Aquatic Organisms 130:51–63.

16. De Los Santos, C. B., D. Krause-Jensen, T. Alcoverro, N. Marbà, C. M. Duarte, M. M. Van Katwijk, M. Pérez, J. Romero, J. L. Sánchez-Lizaso, G. Roca, E. Jankowska, J. L. Pérez-Lloréns, J. Fournier, M. Montefalcone, G. Pergent, J. M. Ruiz, S. Cabaço, K. Cook, R. J. Wilkes, F. E. Moy, G. M.-R. Trayter, X. S. Arañó, D. J. De Jong, Y. Fernández-Torquemada, I. Auby, J. J. Vergara, and R. Santos. 2019. Recent trend reversal for declining European seagrass meadows. Nature Communications 10:3356.

17. Dell, A. I., S. Pawar, and V. M. Savage. 2011. Systematic variation in the temperature dependence of physiological and ecological traits. Proceedings of the National Academy of Sciences 108:10591–10596.

18. Deutsch, C. A., J. J. Tewksbury, R. B. Huey, K. S. Sheldon, C. K. Ghalambor, D. C. Haak, and P. R. Martin. 2008. Impacts of climate warming on terrestrial ectotherms across latitude. Proceedings of the National Academy of Sciences 105:6668–6672.

19. Eigenbrode, S. D., N. A. Bosque-Pérez, and T. S. Davis. 2018. Insect-Borne Plant Pathogens and Their Vectors: Ecology, Evolution, and Complex Interactions. Annual Review of Entomology 63:169–191.

20. Eisenlord, M. E., M. V. Agnew, M. Winningham, O. J. Lobo, A. D. Vompe, B. Wippel, C. S. Friedman, C. D. Harvell, and C. A. Burge. 2024. High infectivity and waterborne transmission of seagrass wasting disease. Royal Society Open Science 11:240663.

21. Elderd, B. D., and J. R. Reilly. 2014. Warmer temperatures increase disease transmission and outbreak intensity in a host–pathogen system. Journal of Animal Ecology 83:838–849.

22. Fox, J., and S. Weisberg. 2011. An R companion to applied regression. Second edition. Sage Publications, Thousand Oaks, CA.

23. Gallana, M., M.-P. Ryser-Degiorgis, T. Wahli, and H. Segner. 2013. Climate change and infectious diseases of wildlife: Altered interactions between pathogens, vectors and hosts. Current Zoology 59:427–437.

24. Gehman, A.-L. M., R. J. Hall, and J. E. Byers. 2018. Host and parasite thermal ecology jointly determine the effect of climate warming on epidemic dynamics. Proceedings of the National Academy of Sciences 115:744–749.

25. Godet, L., J. Fournier, M. van Katwijk, F. Olivier, P. Le Mao, and C. Retière. 2008. Before and after wasting disease in common eelgrass Zostera marina along the French Atlantic coasts: a general overview and first accurate mapping. Diseases of Aquatic Organisms 79:249–255.

26. Gossner, M. M., L. Beenken, K. Arend, D. Begerow, and D. Peršoh. 2021. Insect herbivory facilitates the establishment of an invasive plant pathogen. ISME Communications 1:6.

27. Govers, L. L., W. Suykerbuyk, J. H. T. Hoppenreijs, K. Giesen, T. J. Bouma, and M. M. Van Katwijk. 2015. Rhizome starch as indicator for temperate seagrass winter survival. Ecological Indicators 49:53–60.

28. Graham, O. J., L. R. Aoki, C. A. Burge, and C. D. Harvell. 2024. Invertebrate herbivores influence seagrass wasting disease dynamics. Ecology:e4493.

29. Graham, O. J., L. R. Aoki, T. Stephens, J. Stokes, S. Dayal, B. Rappazzo, C. P. Gomes, and C. D. Harvell. 2021. Effects of Seagrass Wasting Disease on Eelgrass Growth and Belowground Sugar in Natural Meadows. Frontiers in Marine Science 8:768668.

30. Graham, O. J., T. Stephens, B. Rappazzo, C. Klohmann, S. Dayal, E. M. Adamczyk, A. Olson, M. Hessing-Lewis, M. Eisenlord, B. Yang, C. Burge, C. P. Gomes, and D. Harvell. 2023. Deeper habitats and cooler temperatures moderate a climate-driven seagrass disease. Philosophical Transactions of the Royal Society B: Biological Sciences 378:20220016.

31. Greenspan, S. E., D. S. Bower, E. A. Roznik, D. A. Pike, G. Marantelli, R. A. Alford, L. Schwarzkopf, and B. R. Scheffers. 2017. Infection increases vulnerability to climate change via effects on host thermal tolerance. Scientific Reports 7:9349.

32. Groner, M., M. Eisenlord, R. Yoshioka, E. Fiorenza, P. Dawkins, O. Graham, M. Winningham, A. Vompe, N. Rivlin, B. Yang, C. Burge, B. Rappazzo, C. Gomes, and C. Harvell. 2021. Warming sea surface temperatures fuel summer epidemics of eelgrass wasting disease. Marine Ecology Progress Series 679:47–58.

33. Hamann, E., C. Blevins, S. J. Franks, M. I. Jameel, and J. T. Anderson. 2021. Climate change alters plant–herbivore interactions. New Phytologist 229:1894–1910.

34. Hartig, F. 2026. DHARMa: Residual Diagnostics for Hierarchical (Multi-Level / Mixed) Regression Models.

35. Heinze, G., M. Ploner, L. Jiricka, and G. Steiner. 2025. logistf: Firth’s Bias-Reduced Logistic Regression.

36. Heinze, G., and M. Schemper. 2002. A solution to the problem of separation in logistic regression. Statistics in Medicine 21:2409–2419.

37. Hernán, G., M. J. Ortega, A. M. Gándara, I. Castejón, J. Terrados, and F. Tomas. 2017. Future warmer seas: increased stress and susceptibility to grazing in seedlings of a marine habitat-forming species. Global Change Biology 23:4530–4543.

38. Jakobsson-Thor, S., G. Toth, J. Brakel, A. Bockelmann, and H. Pavia. 2018. Seagrass wasting disease varies with salinity and depth in natural Zostera marina populations. Marine Ecology Progress Series 587:105–115.

39. Kimes, N. E., C. J. Grim, W. R. Johnson, N. A. Hasan, B. D. Tall, M. H. Kothary, H. Kiss, A. C. Munk, R. Tapia, L. Green, C. Detter, D. C. Bruce, T. S. Brettin, R. R. Colwell, and P. J. Morris. 2012. Temperature regulation of virulence factors in the pathogen *Vibrio coralliilyticus*. The ISME Journal 6:835–846.

40. King, N. G., N. J. McKeown, D. A. Smale, and P. J. Moore. 2018. The importance of phenotypic plasticity and local adaptation in driving intraspecific variability in thermal niches of marine macrophytes. Ecography 41:1469–1484.

41. Kingsolver, J. G., H. Arthur Woods, L. B. Buckley, K. A. Potter, H. J. MacLean, and J. K. Higgins. 2011. Complex Life Cycles and the Responses of Insects to Climate Change. Integrative and Comparative Biology 51:719–732.

42. Kontopoulos, D.-G., A. Sentis, M. Daufresne, N. Glazman, A. I. Dell, and S. Pawar. 2024. No universal mathematical model for thermal performance curves across traits and taxonomic groups. Nature Communications 15:8855.

43. Lactin, D. J., N. J. Holliday, D. L. Johnson, and R. Craigen. 1995. Improved Rate Model of Temperature-Dependent Development by Arthropods. Environmental Entomology 24:68–75.

44. Lee, K.-S., S. R. Park, and J.-B. Kim. 2005. Production dynamics of the eelgrass, Zostera marina in two bay systems on the south coast of the Korean peninsula. Marine Biology 147:1091–1108.

45. Lefcheck, J. S., D. J. Wilcox, R. R. Murphy, S. R. Marion, and R. J. Orth. 2017. Multiple stressors threaten the imperiled coastal foundation species eelgrass (Zostera marina) in Chesapeake Bay, USA. Global Change Biology 23:3474–3483.

46. Lemoine, N. P., W. A. Drews, D. E. Burkepile, and J. D. Parker. 2013. Increased temperature alters feeding behavior of a generalist herbivore. Oikos 122:1669–1678.

47. López-Urrutia, Á., E. San Martin, R. P. Harris, and X. Irigoien. 2006. Scaling the metabolic balance of the oceans. Proceedings of the National Academy of Sciences 103:8739–8744.

48. Low-Décarie, E., T. G. Boatman, N. Bennett, W. Passfield, A. Gavalás-Olea, P. Siegel, and R. J. Geider. 2017. Predictions of response to temperature are contingent on model choice and data quality. Ecology and Evolution 7:10467–10481.

49. Marsh, J. A., W. C. Dennison, and R. S. Alberte. 1986. Effects of temperature on photosynthesis and respiration in eelgrass (Zostera marina L.). Journal of Experimental Marine Biology and Ecology 101:257–267.

50. Martínez-De León, G., and M. P. Thakur. 2024. Ecological debts induced by heat extremes. Trends in Ecology & Evolution 39:1024–1034.

51. McCann, K., A. Hastings, and G. R. Huxel. 1998. Weak trophic interactions and the balance of nature. Nature.

52. McKone, K., and C. Tanner. 2009. Role of salinity in the susceptibility of eelgrass Zostera marina to the wasting disease pathogen Labyrinthula zosterae. Marine Ecology Progress Series 377:123–130.

53. Mertens, N. L., B. D. Russell, and S. D. Connell. 2015. Escaping herbivory: ocean warming as a refuge for primary producers where consumer metabolism and consumption cannot pursue. Oecologia 179:1223–1229.

54. Moffitt, J., and C. Cottam. 1941. Eelgrass depletion on the Pacific coast and its effect upon black brant. Page 26. Report, Washington, D.C.

55. Mordecai, E. A., J. M. Caldwell, M. K. Grossman, C. A. Lippi, L. R. Johnson, M. Neira, J. R. Rohr, S. J. Ryan, V. Savage, M. S. Shocket, R. Sippy, A. M. Stewart Ibarra, M. B. Thomas, and O. Villena. 2019. Thermal biology of mosquito-borne disease. Ecology Letters 71:411.

56. Moreno-Marín, F., F. G. Brun, and M. F. Pedersen. 2018. Additive response to multiple environmental stressors in the seagrass *Zostera marina* L. Limnology and Oceanography 63:1528–1544.

57. Muehlstein, L. 1989. Perspectives on the wasting disease of eelgrass Zostera marina. Diseases of Aquatic Organisms 7:211–221.

58. Murray, N. A., K. DuBois, and J. J. Stachowicz. 2024. Herbivores can benefit both plants and their pathogens through selective herbivory on diseased tissue. Journal of Ecology 112:1413–1424.

59. Nejrup, L. B., and M. F. Pedersen. 2008. Effects of salinity and water temperature on the ecological performance of Zostera marina. Aquatic Botany 88:239–246.

60. Nicolet, K. J., K. M. Chong-Seng, M. S. Pratchett, B. L. Willis, and M. O. Hoogenboom. 2018. Predation scars may influence host susceptibility to pathogens: evaluating the role of corallivores as vectors of coral disease. Scientific Reports 8:5258.

61. Niu, S., P. Zhang, J. Liu, D. Guo, and X. Zhang. 2012. The effect of temperature on the survival, growth, photosynthesis, and respiration of young seedlings of eelgrass Zostera marina L. Aquaculture 350–353:98–108.

62. O’Connor, M. I. 2009. Warming strengthens an herbivore-plant interaction. Ecology:388–398.

63. Orth, R. J., T. J. B. Carruthers, W. C. Dennison, C. M. Duarte, J. W. Fourqurean, K. L. Heck, A. R. Hughes, G. A. Kendrick, W. J. Kenworthy, S. Olyarnik, F. T. Short, M. Waycott, and S. L. Williams. 2006. A Global Crisis for Seagrass Ecosystems. BioScience 56:987.

64. Osenberg, C. W., O. Sarnelle, and S. D. Cooper. 1997. Effect Size in Ecological Experiments: The Application of Biological Models in Meta-Analysis. The American Naturalist 150:798–812.

65. Paaijmans, K. P., S. Blanford, B. H. K. Chan, and M. B. Thomas. 2012. Warmer temperatures reduce the vectorial capacity of malaria mosquitoes. Biology Letters 8:465–468.

66. Padfield, D., M. Castledine, and A. Buckling. 2020. Temperature-dependent changes to host–parasite interactions alter the thermal performance of a bacterial host. The ISME Journal 14:389–398.

67. Padfield, D., and H. O’Sullivan. 2023. rTPC: Fitting and Analysing Thermal Performance Curves.

68. Pagán, I., and F. García-Arenal. 2020. Tolerance of Plants to Pathogens: A Unifying View. Annual Review of Phytopathology 58:77–96.

69. Pincebourde, S., and J. Casas. 2019. Narrow safety margin in the phyllosphere during thermal extremes. Proceedings of the National Academy of Sciences 116:5588–5596.

70. R Core Team. 2025. R: A Language and Environment for Statistical Computing. R Foundation for Statistical Computing, Vienna, Austria.

71. Ralph, P., and F. Short. 2002. Impact of the wasting disease pathogen, Labyrinthula zosterae, on the photobiology of eelgrass Zostera marina. Marine Ecology Progress Series 226:265–271.

72. Rappazzo, B. H., M. E. Eisenlord, O. J. Graham, L. R. Aoki, P. D. Dawkins, D. Harvell, and C. Gomes. 2021. EeLISA: Combating Global Warming Through the Rapid Analysis of Eelgrass Wasting Disease. Proceedings of the AAAI Conference on Artificial Intelligence 35:15156–15165.

73. Rebolledo, A. P., C. M. Sgrò, and K. Monro. 2021. Thermal Performance Curves Are Shaped by Prior Thermal Environment in Early Life. Frontiers in Physiology 12:738338.

74. Reynolds, L. K., K. DuBois, J. M. Abbott, S. L. Williams, and J. J. Stachowicz. 2016. Response of a Habitat-Forming Marine Plant to a Simulated Warming Event Is Delayed, Genotype Specific, and Varies with Phenology. PLOS ONE 11:e0154532.

75. Sanford, E., J. L. Sones, M. García-Reyes, J. H. R. Goddard, and J. L. Largier. 2019. Widespread shifts in the coastal biota of northern California during the 2014–2016 marine heatwaves. Scientific Reports 9:4216.

76. Schenck, F. R., K. DuBois, M. R. Kardish, J. J. Stachowicz, and A. R. Hughes. 2023. The effect of warming on seagrass wasting disease depends on host genotypic identity and diversity. Ecology 104:e3959.

77. Schiebelhut, L. M., R. K. Grosberg, J. J. Stachowicz, and R. A. Bay. 2023. Genomic responses to parallel temperature gradients in the eelgrass *Zostera marina* in adjacent bays. Molecular Ecology 32:2835–2849.

78. Schneider, C. A., W. S. Rasband, and K. W. Eliceiri. 2012. NIH Image to ImageJ: 25 years of image analysis. Nature Methods 9:671–675.

79. Shocket, M. S., A. B. Verwillow, M. G. Numazu, H. Slamani, J. M. Cohen, F. El Moustaid, J. Rohr, L. R. Johnson, and E. A. Mordecai. 2020. Transmission of West Nile and five other temperate mosquito-borne viruses peaks at temperatures between 23°C and 26°C. eLife 9:e58511.

80. Short, F. T., L. K. Muehlstein, and D. Porter. 1987. Eelgrass wasting disease: cause and recurrence of a marine epidemic. The Biological Bulletin 173:557–562.

81. Sinclair, B. J., K. E. Marshall, M. A. Sewell, D. L. Levesque, C. S. Willett, S. Slotsbo, Y. Dong, C. D. G. Harley, D. J. Marshall, B. S. Helmuth, and R. B. Huey. 2016. Can we predict ectotherm responses to climate change using thermal performance curves and body temperatures? Ecology Letters 19:1372–1385.

82. Sokolova, I. 2021. Bioenergetics in environmental adaptation and stress tolerance of aquatic ectotherms: linking physiology and ecology in a multi-stressor landscape. Journal of Experimental Biology 224:jeb236802.

83. Stauffer, R. C. 1937. Changes in the Invertebrate Community of a Lagoon After Disapperance of the Eel Grass. Ecology 18:427–431.

84. Teckentrup, L., V. Grimm, S. Kramer-Schadt, and F. Jeltsch. 2018. Community consequences of foraging under fear. Ecological Modelling 383:80–90.

85. Thayer, G., WJ. Kenworthy, and M. S. Fonseca. 1984. The Ecology of Eelgrass Meadows of the Atlantic Coast: A Community Profile. Page 147. U.S. Fish & Wildlife Service.

86. Therneau, T. 2024. A Package for Survival Analysis in R.

87. Therneau, T. M., and P. M. Grambsch. 2000. Modeling Survival Data: Extending the Cox Model. Springer New York, New York, NY.

88. Touchette, B. W., and J. M. Burkholder. 2000. Overview of the physiological ecology of carbon metabolism in seagrasses. Journal of Experimental Marine Biology and Ecology 250:169–205.

89. Trevathan-Tackett, S. M., B. K. Sullivan, K. Robinson, O. Lilje, P. I. Macreadie, and F. H. Gleason. 2018. Pathogenic Labyrinthula associated with Australian seagrasses: Considerations for seagrass wasting disease in the southern hemisphere. Microbiological Research 206:74–81.

90. Tutin, T. G. 1942. Zostera L. Journal of Ecology 30:217–226.

91. Wan, L., G. Liu, H. Cheng, S. Yang, Y. Shen, and X. Su. 2023. Global warming changes biomass and C:N:P stoichiometry of different components in terrestrial ecosystems. Global Change Biology 29:7102–7116.

92. Ware-Gilmore, F., C. M. Sgrò, Z. Xi, H. L. C. Dutra, M. J. Jones, K. Shea, M. D. Hall, M. B. Thomas, and E. A. McGraw. 2021. Microbes increase thermal sensitivity in the mosquito Aedes aegypti, with the potential to change disease distributions. PLOS Neglected Tropical Diseases 15:e0009548.

93. West, D. C., and D. M. Post. 2016. Impacts of warming revealed by linking resource growth rates with consumer functional responses. Journal of Animal Ecology 85:671–680.

94. Westera, M. B., and P. S. Lavery. 2006. A comparison of hole punch and needle punch methods for the measurement of seagrass productivity. Aquatic Botany 85:267–269.

95. Yvon-Durocher, G., J. I. Jones, M. Trimmer, G. Woodward, and J. M. Montoya. 2010. Warming alters the metabolic balance of ecosystems. Philosophical Transactions of the Royal Society B: Biological Sciences 365:2117–2126.

96. Zvereva, E. L., and M. V. Kozlov. 2021. Latitudinal gradient in the intensity of biotic interactions in terrestrial ecosystems: Sources of variation and differences from the diversity gradient revealed by meta-analysis. Ecology Letters 24:2506–2520.

