## Supplemental tables, figures, appendices for "Distinct thermal responses of a host plant and an invertebrate herbivore affect ecosystem productivity and disease dynamics in a coastal marine ecosystem"

**Supplementary Material 1**
for:

**Distinct thermal responses of a host plant and an invertebrate herbivore affect ecosystem productivity and disease dynamics in a coastal marine ecosystem**

**Description**: Supplementary tables, figures, and model equation appendices.

TABLE OF CONTENTS

Figure S1: Photo of rhizome lesions p. 3

Table S1: Summary of plants with a leaf that became detached in mesocosm experiment p. 4

Appendix 1: Equations for the models fit to eelgrass responses in the no-grazing treatments. p. 5

Appendix 2: Equations for the models fit to eelgrass responses in the grazing treatments. p. 6

Appendix 3: Equations for the nonlinear models fit to final disease severity. p. 7

Appendix 4) Mesocosm experiment temperature treatment summary. p. 8

Table S2: Summary of mesocosm experiment treatment temperatures

Figure S2: Summary of mesocosm experiment treatment temperature vs. time

Appendix 5) Grazing experiment summary. p. 9

Table S3: Summary of grazing experiment treatment temperatures

Table S4: Summary of isopod sample size in each temperature treatment

Figure S3: Thermal performance curves for eelgrass above- & belowground dry mass. p. 10

Figure S4: Thermal performance curve $T_{opt}$ & $T_{br}$ estimates for eelgrass responses. p. 11

Figure S5: Thermal performance curve $r_{max}$ estimates for eelgrass responses. p. 12

Table S5: Thermal performance curve derived parameter estimates ($T_{opt}$, $T_{br}$, $r_{max}$) for eelgrass responses (no-grazing treatments). p. 13

Table S6: Thermal performance curve parameter estimates (a, b, etc.) for eelgrass responses (no-grazing treatments). p. 14-16

Table S7: Model parameter estimates for eelgrass responses in the grazing treatments. p. 17-18

Figure S6: Interaction strength vs. temperature in the mesocosm experiment. p. 19

Figure S7: Rhizome necroses vs. temperature in the mesocosm experiment. p. 20

Figure S8: Final disease severity vs. temperature. p. 21

Figure S9: Final diseased area vs. temperature – pooled data fit. p. 21

Table S8: Parameter estimates for exponential models fit to final disease area and severity. p. 22
Table S9: Statistical results for Gamma GLM fit to final diseased area. p. 23
Table S10: Derived parameter estimates for isopod thermal performance curves. p. 24
Table S11: Model parameter estimates for isopod thermal performance curves. p. 24
Figure S10: Hypothetical response of top-down interaction strength (effect of herbivores on net productivity) vs. temperature under scenarios with herbivores with contrasting thermal tolerance. p. 25

**Figure S1)** Photo of rhizome necroses/damage that were noted (presence/absence) at the end of the experiment. Experimental plants did not exhibit these darkened areas when they were planted at the start of the experiment.

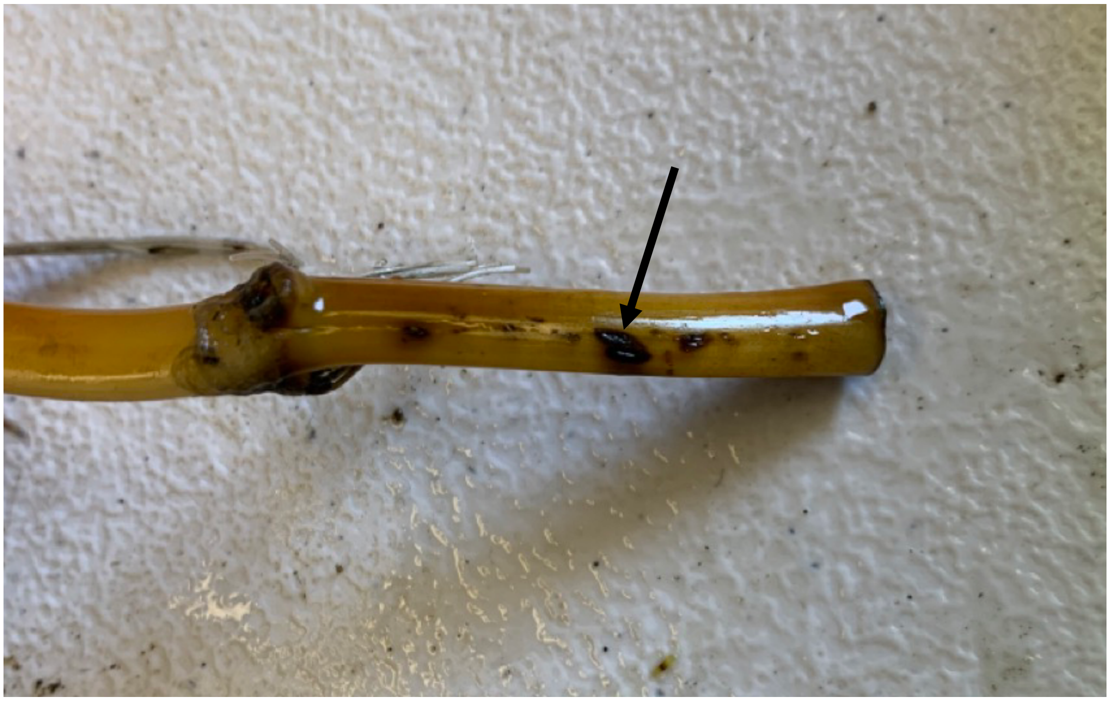

**Table S1)** Summary of plants with a detached leaf or partial leaf at the end of the experiment. An asterisk (*) is used to indicate the number of plants that were considered dead in a treatment at the end of the experiment.

| Treatment | Temp trt. | No. of plants |
|---|---|---|
| healthy-no grazing | 19.3 | 1 |
| healthy-grazing | 19.3 | 1 |
| diseased-no grazing | 19.3 | 1 |
| diseased-grazing | 22.6 | 1 |
| healthy-grazing | 22.6 | 1 |
| diseased-grazing | 24.8 | 1* |
| diseased-no grazing | 24.8 | 1* |

**Appendix 1)** Equations for the TPC models fit to eelgrass responses in the **no-grazing** treatments in the mesocosm experiment, and isopod responses (mesocosm experiment and grazing experiment).

1) Lactin 2 model:

$$rate = exp^{aT} - exp^{aT_{max} - (\frac{T_{max} - T}{\delta_T})} + b$$

2) Weibull model

$$rate = a\left(\frac{c-1}{c}\right)^{1-c/c}\left(\frac{T-T_{opt}}{b} + \left(\frac{c-1}{c}\right)^{1/c}\right)^{c-1} exp^{-\left(\frac{T-T_{opt}}{b} + \left(\frac{c-1}{c}\right)^{1/c}\right)^{c}} + \frac{c-1}{c}$$

3) 2nd-order polynomial (quadratic) model:

$$rate = a + bT + cT^2$$

4) Binomial regression with 2nd-order polynomial (quadratic) effect of temperature:

$$logit(probability) = a + bT + cT^2$$

**Appendix 2)** Equations for the models fit to eelgrass responses in the **grazing** treatments in the mesocosm experiment.

1) Linear model

$$rate = a + bT$$

2) Power model

$$rate = c + aT^b$$

3) Logarithmic model

$$rate = a + b \cdot ln(T)$$

4) Asymptotic regression model

$$rate = a - (a - b)exp^{(-cT)}$$

**Appendix 3)** Equations for the models fit to final diseased area and disease severity in the mesocosm experiment

1) Linear model

$rate = a + bT$

2) Exponential model

$rate = a \cdot exp^{bT}$

3) Power model:

$rate = c + aT^{b}$

4) 2nd order polynomial (quadratic) model

$rate = a + bT + cT^{2}$

**Appendix 4) Mesocosm experiment temperature treatment summary**

**Table S2)** Summary of the mean temperatures in each treatment in the mesocosm experiment.

| Temp. trt. | No. timepoints | Mean temp. (°C) | SE |
|---|---|---|---|
| 1 | 1,633 | 8.95 | 0.02 |
| 2 | 1,633 | 15.80 | 0.02 |
| 3 | 1,633 | 19.31 | 0.01 |
| 4 | 1,632 | 22.56 | 0.01 |
| 5 | 1,632 | 24.78 | 0.01 |

**Figure S2)** Water temperatures for each temperature treatment (indicated by color) over the duration of the mesocosm experiment. Points show the mean of two water tables per treatment at each 15-minute interval. Error bars (in black) indicate ± SE. Error bars are not visible at most time points, reflecting minimal temperature variation between water tables within a treatment.

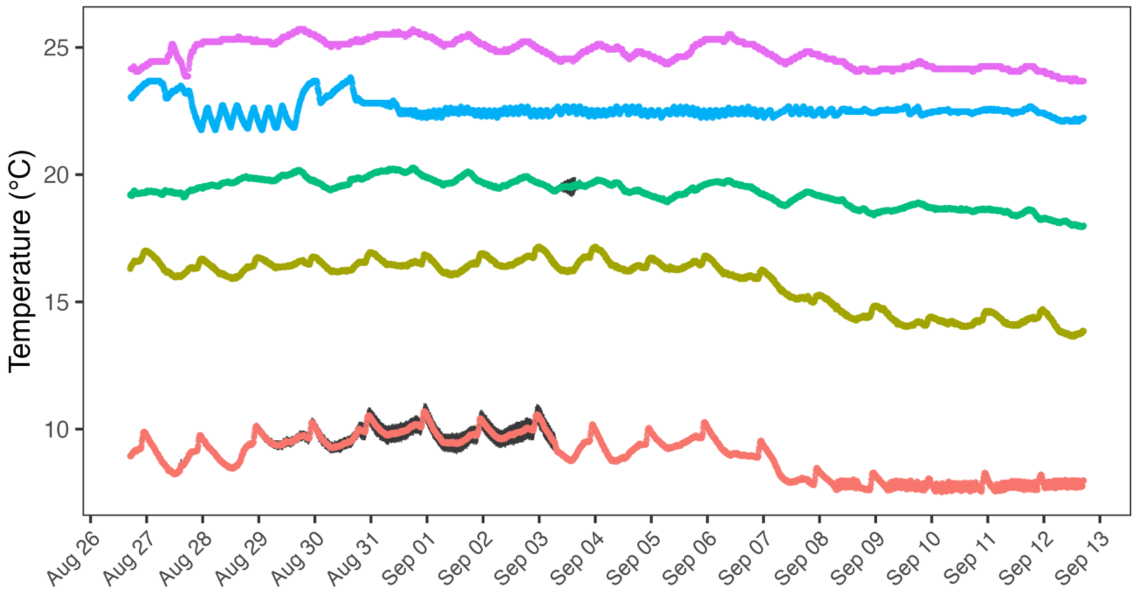

**Appendix 5) Grazing experiment summary**

**Table S3)** Summary of the mean temperatures in each treatment in the isopod grazing experiment.

| Temp. trt. | No. timepoints | Mean temp. (°C) | SE |
|---|---|---|---|
| 1 | 1,009 | 8.42 | 0.02 |
| 2 | 1,009 | 13.41 | 0.04 |
| 3 | 1,009 | 15.12 | 0.03 |
| 4 | 1,009 | 18.88 | 0.02 |
| 5 | 1,009 | 22.42 | 0.01 |
| 6 | 1,009 | 24.41 | 0.01 |

**Table S4)** Summary of the number of isopods in each temperature treatment in the isopod grazing experiment. Each temperature treatment started with 16 isopods, but individuals that died or escaped during the trials were excluded from analyses.

| Temp. trt. | Mean temp. (°C) | n |
|---|---|---|
| 1 | 8.4 | 16 |
| 2 | 13.4 | 15 |
| 3 | 15.1 | 16 |
| 4 | 18.9 | 15 |
| 5 | 22.4 | 14 |
| 6 | 24.4 | 11 |

**Figure S3)** Belowground **(a)** vs. aboveground **(b)** final dry mass (g) and **(c)** final root:shoot ratio (belowground: aboveground dry mass). Colors indicate disease exposure and grazing treatments. Lines show the best-fit predictions for a second-order polynomial model, with shaded ribbons indicating 95% CI.  Points show individual focal plant replicates.

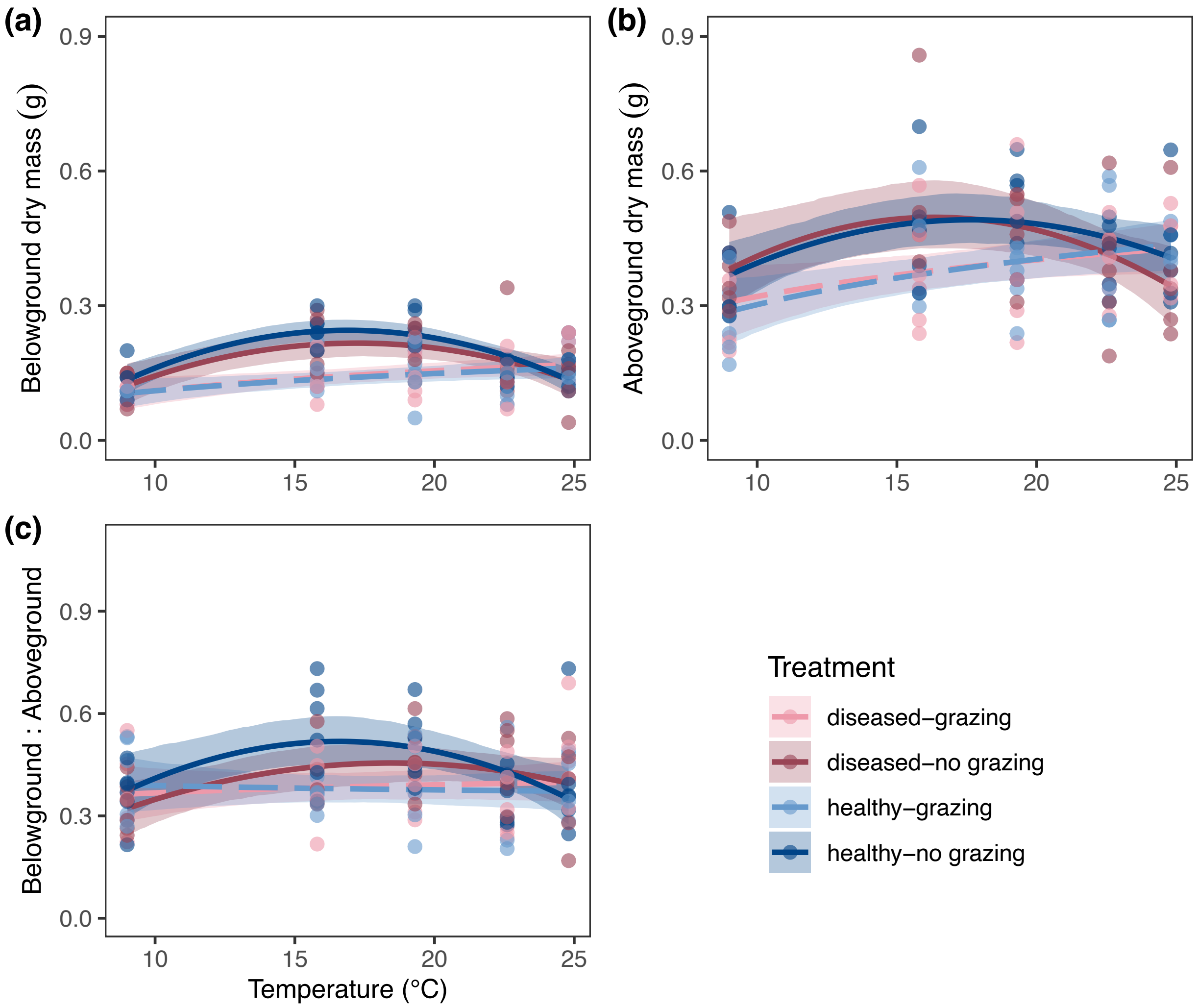

**Figure S4)** Summary of TPC model parameter estimates and 95% CI for different plant responses in the mesocosm experiment. Note, for all plant responses except rhizome lesions only the no grazing treatments were used.

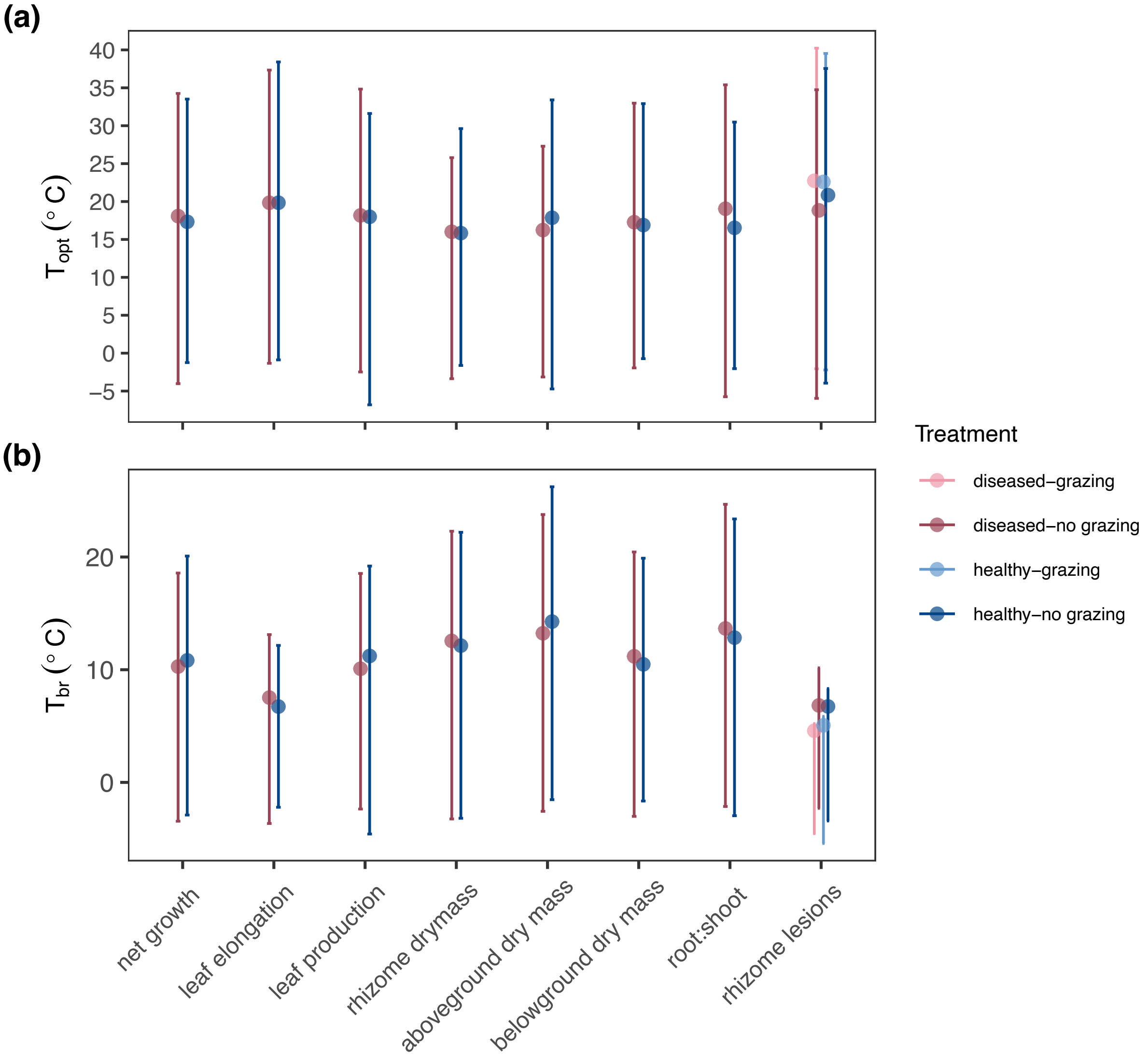

**Figure S5)** Summary of TPC model estimates and 95% CI for $r_{max}$ of each plant response in each no-grazing, disease exposure treatment combination (healthy-no grazing and diseased-no grazing) in the mesocosm experiment.

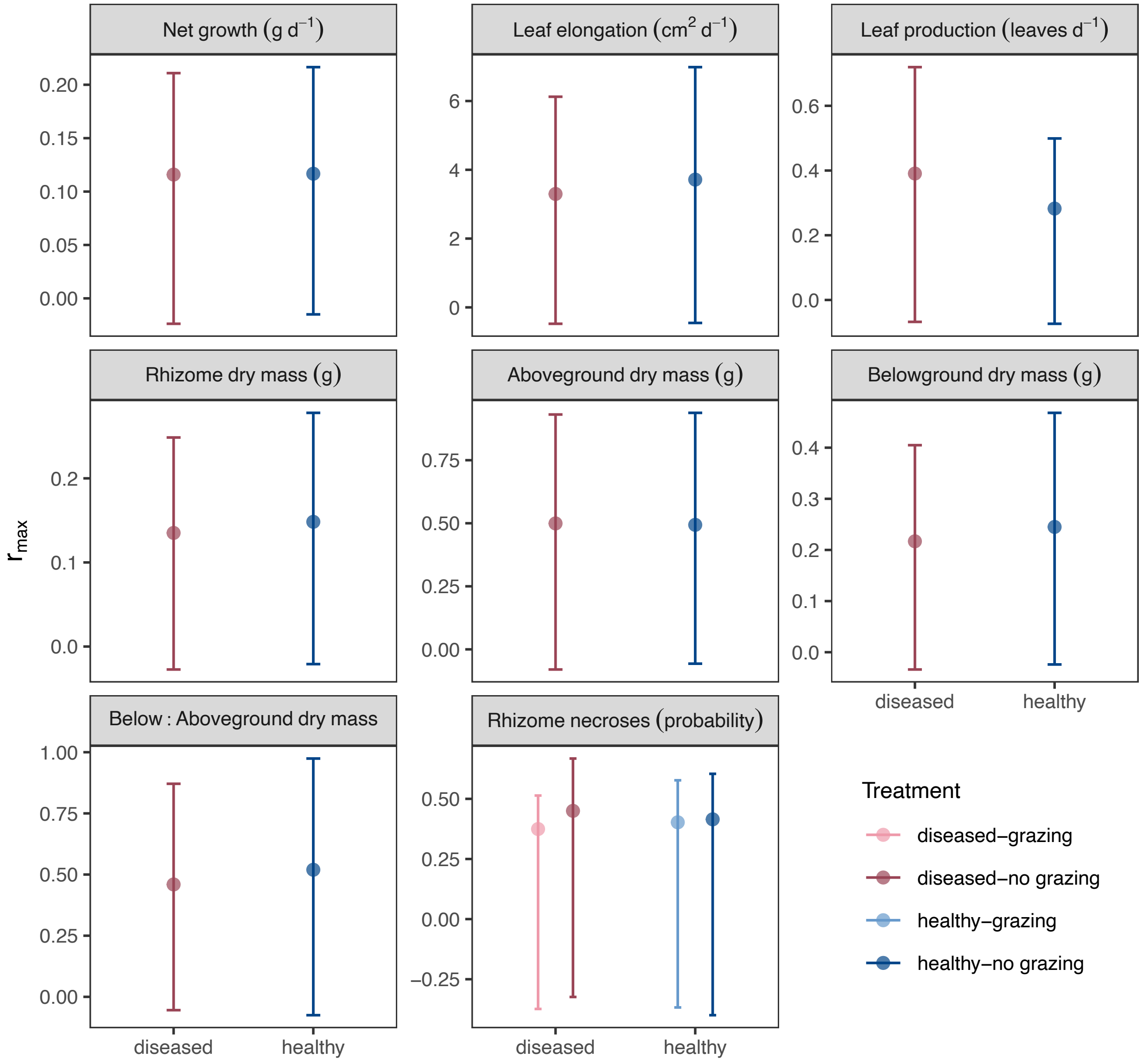

**Table S5)** Derived parameter estimates and 95% CI for TPCs fit to each eelgrass response in the mesocosm experiment. (Parameter estimates and error are also visualized in Figures S4-S5).

| | | | | Thermal optimum (°C) | | Maximum performance | | Thermal Breadth (°C) | |
|---|---|---|---|---|---|---|---|---|---|
| **Response** | **Disease treatment** | **Grazing treatment** | **Model** | **$T_{opt}$** | **95% CI** | **$r_{max}$** | **95% CI** | **$T_{br}$** | **95% CI** |
| net growth | healthy | no grazing | poly2 | 17.3 | [16.2, 18.6] | 0.1 | [0.1, 0.1] | 10.8 | [9.3, 13.7] |
| net growth | diseased | no grazing | poly2 | 18.1 | [16.2, 22.1] | 0.1 | [0.1, 0.1] | 10.3 | [8.3, 13.7] |
| leaf elongation | healthy | no grazing | Weibull | 19.8 | [18.6, 20.7] | 3.7 | [3.3, 4.2] | 6.7 | [5.4, 8.9] |
| leaf elongation | diseased | no grazing | Weibull | 19.8 | [17.5, 21.2] | 3.3 | [2.8, 3.8] | 7.5 | [5.6, 11.2] |
| leaf production | healthy | no grazing | poly2 | 18.0 | [13.6, 24.8] | 0.3 | [0.2, 0.4] | 11.2 | [8, 15.8] |
| leaf production | diseased | no grazing | poly2 | 18.2 | [16.7, 20.7] | 0.4 | [0.3, 0.5] | 10.1 | [8.5, 12.4] |
| rhizome drymass | healthy | no grazing | poly2 | 15.8 | [13.8, 17.5] | 0.1 | [0.1, 0.2] | 12.1 | [10.1, 15.3] |
| rhizome drymass | diseased | no grazing | poly2 | 16.0 | [9.8, 19.4] | 0.1 | [0.1, 0.2] | 12.6 | [9.7, 15.8] |
| aboveground dry mass | healthy | no grazing | poly2 | 17.9 | [15.5, 22.6] | 0.5 | [0.4, 0.6] | 14.3 | [12, 15.8] |
| aboveground dry mass | diseased | no grazing | poly2 | 16.2 | [11.1, 19.4] | 0.5 | [0.4, 0.6] | 13.2 | [10.5, 15.8] |
| belowground dry mass | healthy | no grazing | poly2 | 16.9 | [16, 17.6] | 0.2 | [0.2, 0.3] | 10.5 | [9.4, 12.1] |
| belowground dry mass | diseased | no grazing | poly2 | 17.3 | [15.7, 19.2] | 0.2 | [0.2, 0.3] | 11.2 | [9.3, 14.2] |
| belowground: aboveground | healthy | no grazing | poly2 | 16.5 | [13.9, 18.6] | 0.5 | [0.5, 0.6] | 12.8 | [10.5, 15.8] |
| belowground: aboveground | diseased | no grazing | poly2 | 19.1 | [16.3, 24.8] | 0.5 | [0.4, 0.5] | 13.7 | [11, 15.8] |
| rhizome lesions | diseased | grazing | poly2 | 22.7 | [17.5, 24.8] | 0.4 | [0.1, 0.7] | 4.6 | [0.6, 9.1] |
| rhizome lesions | healthy | grazing | poly2 | 22.6 | [16.9, 24.8] | 0.4 | [0.2, 0.8] | 5.1 | [0.8, 10.5] |
| rhizome lesions | healthy | no grazing | poly2 | 20.8 | [16.7, 24.8] | 0.4 | [0.2, 0.8] | 6.7 | [1.6, 10.2] |
| rhizome lesions | diseased | no grazing | poly2 | 18.8 | [15.9, 24.8] | 0.5 | [0.2, 0.8] | 6.8 | [3.3, 9.1] |

**Table S6)** Parameter estimates for TPC models fit to each eelgrass response (no-grazing treatments) in the mesocosm experiment. Equations with parameters for each model are described in Appendix 1.

| Response | Disease trt. | Grazing trt. | Model | Parameter | Estimate | 95% CI |
|---|---|---|---|---|---|---|
| net growth | healthy | no grazing | poly2 | a | -0.1 | [-0.2, 0] |
| net growth | healthy | no grazing | poly2 | b | 0.0 | [0, 0] |
| net growth | healthy | no grazing | poly2 | c | 0.0 | [0, 0] |
| net growth | diseased | no grazing | poly2 | a | -0.2 | [-0.3, 0] |
| net growth | diseased | no grazing | poly2 | b | 0.0 | [0, 0] |
| net growth | diseased | no grazing | poly2 | c | 0.0 | [0, 0] |
| leaf elongation | healthy | no grazing | Weibull | a | 3.7 | [3.3, 4.2] |
| leaf elongation | healthy | no grazing | Weibull | topt | 19.8 | [18.6, 20.7] |
| leaf elongation | healthy | no grazing | Weibull | b | 18,518.7 | [14.4, 139417.8] |
| leaf elongation | healthy | no grazing | Weibull | c | 3,812.5 | [2.5, 29216.5] |
| leaf elongation | diseased | no grazing | Weibull | a | 3.3 | [2.8, 3.8] |
| leaf elongation | diseased | no grazing | Weibull | topt | 19.8 | [17.5, 21.2] |
| leaf elongation | diseased | no grazing | Weibull | b | 21,717.7 | [14.3, 147357] |
| leaf elongation | diseased | no grazing | Weibull | c | 4,227.1 | [1.7, 30575.8] |
| leaf production | healthy | no grazing | poly2 | a | -0.2 | [-0.7, 0.3] |
| leaf production | healthy | no grazing | poly2 | b | 0.1 | [0, 0.1] |
| leaf production | healthy | no grazing | poly2 | c | 0.0 | [0, 0] |
| leaf production | diseased | no grazing | poly2 | a | -0.6 | [-1, -0.2] |
| leaf production | diseased | no grazing | poly2 | b | 0.1 | [0.1, 0.2] |
| leaf production | diseased | no grazing | poly2 | c | 0.0 | [0, 0] |
| rhizome dry mass | healthy | no grazing | poly2 | a | -0.1 | [-0.2, 0.1] |
| rhizome dry mass | healthy | no grazing | poly2 | b | 0.0 | [0, 0] |
| rhizome dry mass | healthy | no grazing | poly2 | c | 0.0 | [0, 0] |
| rhizome dry mass | diseased | no grazing | poly2 | a | 0.0 | [-0.2, 0.1] |
| rhizome dry mass | diseased | no grazing | poly2 | b | 0.0 | [0, 0] |

| Response | Disease trt. | Grazing trt. | Model | Parameter | Estimate | 95% CI |
|---|---|---|---|---|---|---|
| rhizome dry mass | diseased | no grazing | poly2 | c | 0.0 | [0, 0] |
| belowground dry mass | healthy | no grazing | poly2 | a | -0.3 | [-0.4, -0.1] |
| belowground dry mass | healthy | no grazing | poly2 | b | 0.1 | [0, 0.1] |
| belowground dry mass | healthy | no grazing | poly2 | c | 0.0 | [0, 0] |
| belowground dry mass | diseased | no grazing | poly2 | a | -0.2 | [-0.4, 0] |
| belowground dry mass | diseased | no grazing | poly2 | b | 0.0 | [0, 0.1] |
| belowground dry mass | diseased | no grazing | poly2 | c | 0.0 | [0, 0] |
| aboveground dry mass | healthy | no grazing | poly2 | a | 0.0 | [-0.4, 0.3] |
| aboveground dry mass | healthy | no grazing | poly2 | b | 0.1 | [0, 0.1] |
| aboveground dry mass | healthy | no grazing | poly2 | c | 0.0 | [0, 0] |
| aboveground dry mass | diseased | no grazing | poly2 | a | -0.1 | [-0.5, 0.4] |
| aboveground dry mass | diseased | no grazing | poly2 | b | 0.1 | [0, 0.1] |
| aboveground dry mass | diseased | no grazing | poly2 | c | 0.0 | [0, 0] |
| belowground: aboveground | healthy | no grazing | poly2 | a | -0.2 | [-0.6, 0.3] |
| belowground: aboveground | healthy | no grazing | poly2 | b | 0.1 | [0, 0.1] |
| belowground: aboveground | healthy | no grazing | poly2 | c | 0.0 | [0, 0] |
| belowground: aboveground | diseased | no grazing | poly2 | a | -0.1 | [-0.4, 0.3] |
| belowground: aboveground | diseased | no grazing | poly2 | b | 0.1 | [0, 0.1] |

| Response | Disease trt. | Grazing trt. | Model | Parameter | Estimate | 95% CI |
|---|---|---|---|---|---|---|
| belowground: aboveground | diseased | no grazing | poly2 | c | 0.0 | [0, 0] |
| rhizome necroses | healthy | no grazing | binomial-poly2 | a | -13.0 | [-49.2, -2.3] |
| rhizome necroses | healthy | no grazing | binomial-poly2 | b | 1.2 | [-0.1, 5.1] |
| rhizome necroses | healthy | no grazing | binomial-poly2 | c | 0.0 | [-0.1, 0] |
| rhizome necroses | diseased | no grazing | binomial-poly2 | a | -13.6 | [-63, -4.7] |
| rhizome necroses | diseased | no grazing | binomial-poly2 | b | 1.4 | [0.3, 6.2] |
| rhizome necroses | diseased | no grazing | binomial-poly2 | c | 0.0 | [-0.2, 0] |
| rhizome necroses | diseased | grazing | binomial-poly2 | a | -9.7 | [-68.5, 2.5] |
| rhizome necroses | diseased | grazing | binomial-poly2 | b | 0.7 | [-0.8, 6.8] |
| rhizome necroses | diseased | grazing | binomial-poly2 | c | 0.0 | [-0.2, 0] |
| rhizome necroses | healthy | grazing | binomial-poly2 | a | -7.4 | [-14, 1.7] |
| rhizome necroses | healthy | grazing | binomial-poly2 | b | 0.5 | [-0.7, 1.6] |
| rhizome necroses | healthy | grazing | binomial-poly2 | c | 0.0 | [0, 0] |

**Table S7)** Parameter estimates for models fit to each eelgrass response (grazing treatments) in the mesocosm experiment. Equations with parameters for each model are described in Appendix 1.

| Response | Disease trt. | Grazing trt. | Model | Parameter | Estimate | 95% CI |
|---|---|---|---|---|---|---|
| net growth | diseased | grazing | log | a | -0.2 | [-0.3, -0.1] |
| net growth | diseased | grazing | log | b | 0.1 | [0, 0.1] |
| net growth | healthy | grazing | log | a | 0.0 | [-0.2, 0.1] |
| net growth | healthy | grazing | log | b | 0.0 | [0, 0.1] |
| leaf elongation | diseased | grazing | log | a | -2.3 | [-4.9, 0.6] |
| leaf elongation | diseased | grazing | log | b | 1.5 | [0.6, 2.4] |
| leaf elongation | healthy | grazing | log | a | -1.9 | [-4.5, 0.8] |
| leaf elongation | healthy | grazing | log | b | 1.5 | [0.5, 2.4] |
| leaf production | diseased | grazing | log | a | -0.3 | [-0.7, 0.1] |
| leaf production | diseased | grazing | log | b | 0.2 | [0, 0.3] |
| leaf production | healthy | grazing | log | a | -0.3 | [-0.6, 0.2] |
| leaf production | healthy | grazing | log | b | 0.2 | [0, 0.3] |
| rhizome dry mass | diseased | grazing | log | a | 0.1 | [0, 0.2] |
| rhizome dry mass | diseased | grazing | log | b | 0.0 | [0, 0.1] |
| rhizome dry mass | healthy | grazing | log | a | 0.0 | [-0.1, 0.1] |
| rhizome dry mass | healthy | grazing | log | b | 0.0 | [0, 0.1] |
| aboveground dry mass | diseased | grazing | log | a | 0.0 | [-0.2, 0.4] |
| aboveground dry mass | diseased | grazing | log | b | 0.1 | [0, 0.2] |
| aboveground dry mass | healthy | grazing | log | a | 0.0 | [-0.3, 0.3] |
| aboveground dry mass | healthy | grazing | log | b | 0.1 | [0, 0.2] |
| belowground dry mass | diseased | grazing | log | a | 0.0 | [-0.2, 0.1] |
| belowground dry mass | diseased | grazing | log | b | 0.1 | [0, 0.1] |
| belowground dry mass | healthy | grazing | log | a | 0.0 | [-0.1, 0.1] |
| belowground dry mass | healthy | grazing | log | b | 0.1 | [0, 0.1] |

| Response | Disease trt. | Grazing trt. | Model | Parameter | Estimate | 95% CI |
|---|---|---|---|---|---|---|
| below: aboveground | diseased | grazing | log | a | 0.3 | [0, 0.6] |
| below: aboveground | diseased | grazing | log | b | 0.0 | [-0.1, 0.1] |
| below: aboveground | healthy | grazing | log | a | 0.4 | [0.1, 0.8] |
| below: aboveground | healthy | grazing | log | b | 0.0 | [-0.1, 0.1] |

**Figure S6)** Top-down interaction strength, or the effect of isopod grazers on eelgrass growth, estimated for each temperature treatment as the log-response ratio of the mean final total dry mass of eelgrass in the healthy-no grazed treatment vs. the healthy-grazed treatment, divided by the experimental duration. Positive interaction strength values indicate isopods reduce final eelgrass dry mass, negative values indicate they increase it. Points indicate the mean interaction strength, with error bars showing 95% CI.  The line shows the predictions for the best-fit model (Lactin2).

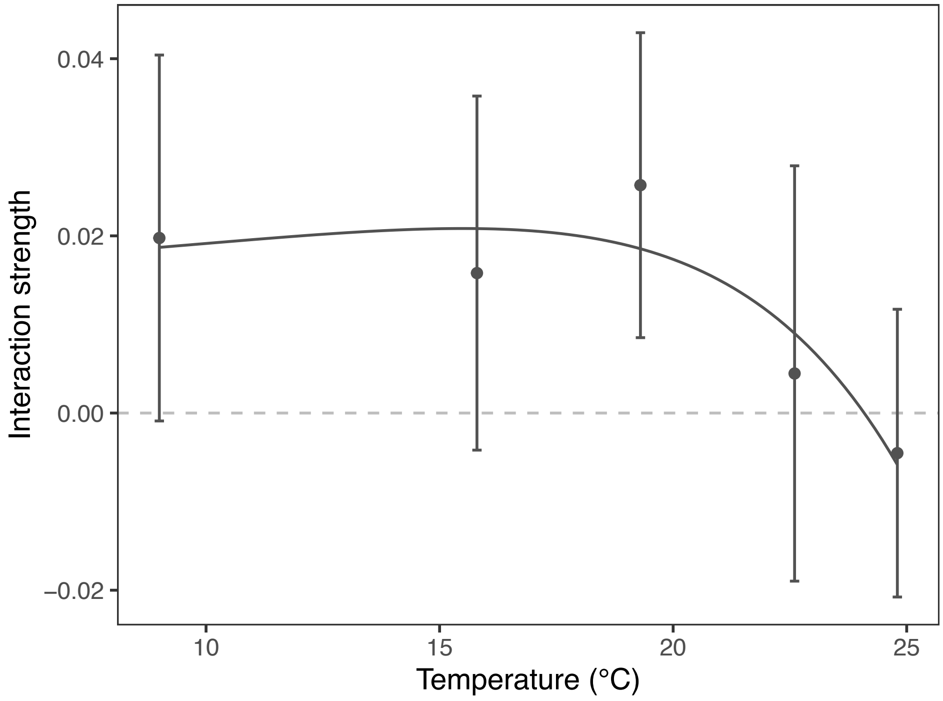

**Figure S7)** Rhizome necroses vs. temperature at the end of the mesocosm experiment. Lines show the best-fit predictions from a binomial regression (logit link) with quadratic effect of temperature, with shaded ribbons indicating the 95% CI. Points indicate the proportion of plants in each treatment group that had rhizome necroses (excluding plants that died). A small horizontal jitter was added to the points so overlapping points could be seen more easily.

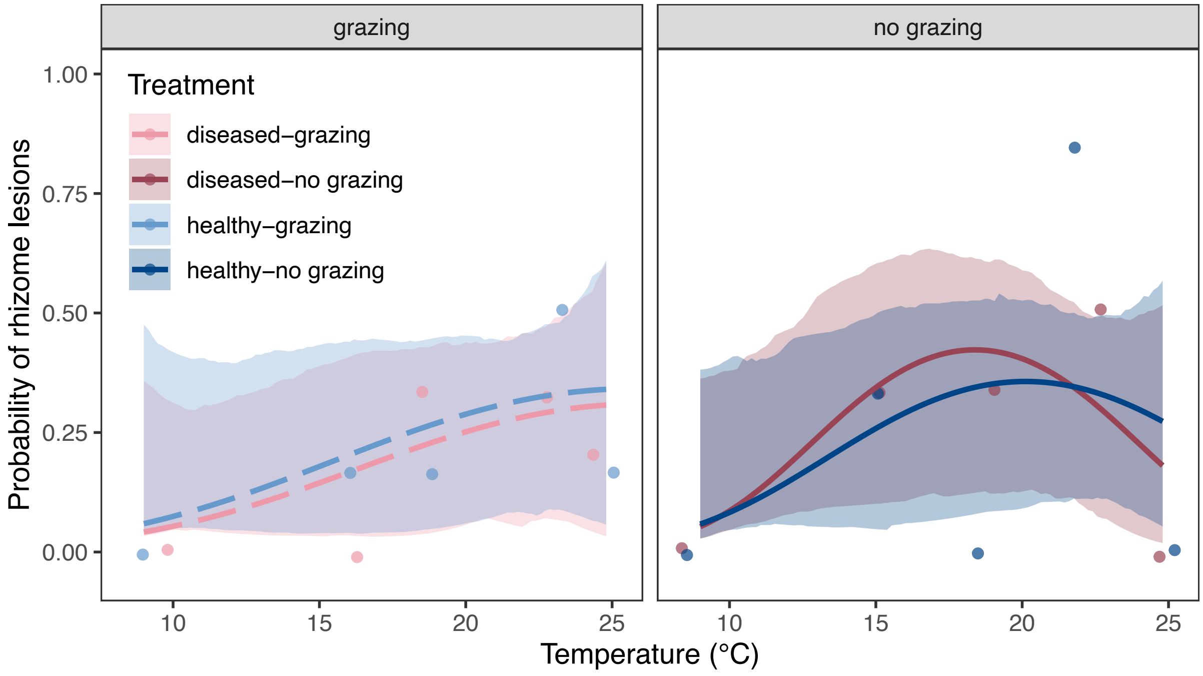

**Figure S8)** Final disease severity (i.e., the proportion of lesioned leaf area out of total leaf area on a plant) vs. temperature in the mesocosm experiment. Lines show best-fit (exponential) model predictions, with 95% CI. Points show observed responses of individual focal plant replicates. Dashed lines indicate grazing treatments, solid lines indicate no-grazing treatments.

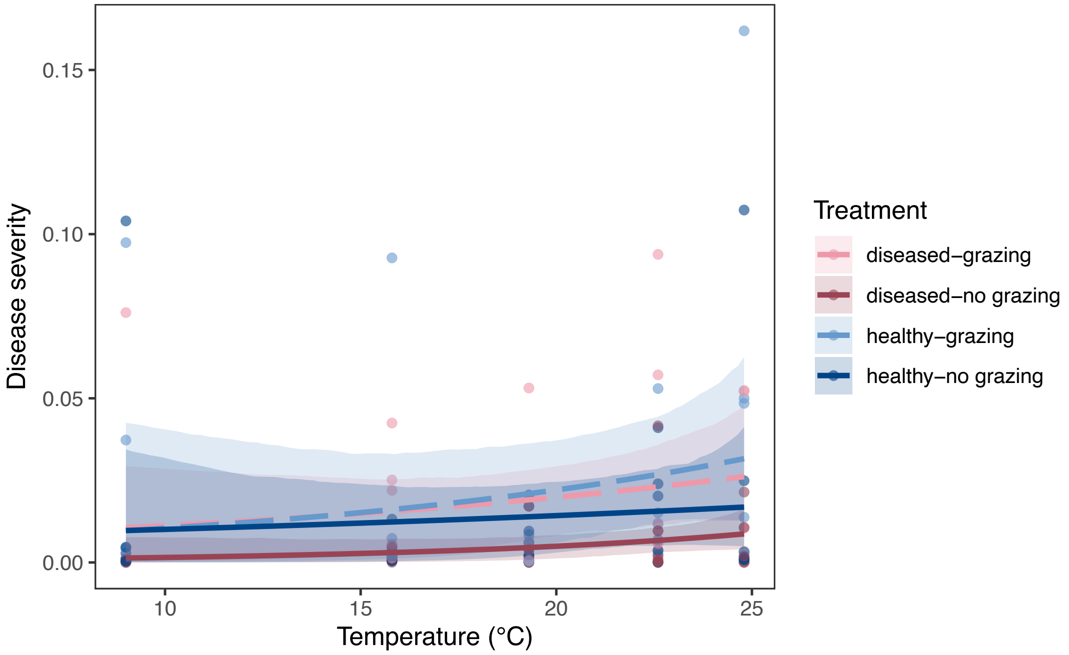

**Figure S9)** Final diseased area ($mm^2$) vs. temperature in the mesocosm experiment, pooled across all grazing and disease exposure treatments. The line and shaded ribbon show exponential model predictions with 95% CI. Points show individual focal plant replicates.

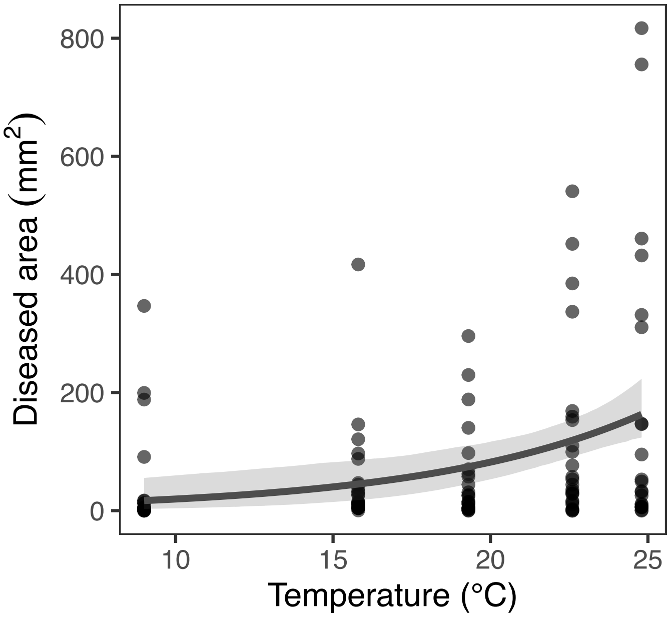

**Table S8)** Parameter estimates for the exponential model ($rate = a \cdot exp^{bT}$) fit to the final disease area and final disease severity data in the mesocosm experiment.

| Response | Disease trt. | Grazing trt. | Model | Parameter | Estimate | 95% CI |
|---|---|---|---|---|---|---|
| disease area | diseased | grazing | exponential | a | 57.88 | [4.73, 128.72] |
| disease area | diseased | grazing | exponential | b | 0.15 | [0.02, 0.42] |
| disease area | healthy | grazing | exponential | a | 19.44 | [0.01, 98.11] |
| disease area | healthy | grazing | exponential | b | 0.43 | [0.08, 1.09] |
| disease area | healthy | no grazing | exponential | a | 55.48 | [1.11, 138.89] |
| disease area | healthy | no grazing | exponential | b | 0.15 | [-0.03, 0.54] |
| disease area | diseased | no grazing | exponential | a | 24.02 | [0.37, 58.14] |
| disease area | diseased | no grazing | exponential | b | 0.13 | [-0.07, 0.58] |
| disease severity | diseased | grazing | exponential | a | 0.01 | [0, 0.03] |
| disease severity | diseased | grazing | exponential | b | 0.08 | [-0.04, 0.3] |
| disease severity | healthy | grazing | exponential | a | 0.02 | [0, 0.03] |
| disease severity | healthy | grazing | exponential | b | 0.10 | [-0.06, 0.4] |
| disease severity | healthy | no grazing | exponential | a | 0.01 | [0, 0.02] |
| disease severity | healthy | no grazing | exponential | b | 0.06 | [-0.1, 0.45] |
| disease severity | diseased | no grazing | exponential | a | 0.00 | [0, 0.01] |
| disease severity | diseased | no grazing | exponential | b | 0.15 | [-0.03, 0.5] |

**Table S9)** Results for the best-fit Gamma GLM (log link) for final diseased area. Statistics are Type II likelihood ratio tests. Significant predictors are highlighted in **bold**.

| Response | Predictor | $\chi^2$ | df | P-value |
|---|---|---|---|---|
| diseased area | Disease treatment | 1.41 | 1 | 0.236 |
| | Grazing treatment | 2.58 | 1 | 0.108 |
| | **Temperature** | **11.61** | **1** | **<0.001** |
| | Disease × Grazing | 1.70 | 1 | 0.192 |

**Table S10)** Derived parameter estimates for the thermal performance curves (TPCs) of isopod responses ($\pm$ 95% CI) for the mesocosm and grazing (isopod only) experiments.

| | | | Thermal optimum (°C) | | Maximum performance | | Thermal Breadth (°C) | |
|---|---|---|---|---|---|---|---|---|
| **Response** | **Disease treatment** | **Model** | $T_{opt}$ | **95% CI** | $r_{max}$ | **95% CI** | $T_{br}$ | **95% CI** |
| isopod survival | diseased | binomial-poly2 | 13.8 | [9, 16.1] | 0.9 | [0.8, 1] | 10.6 | [8.6, 12.5] |
| isopod survival | healthy | binomial-poly2 | 15.4 | [13.8, 16.3] | 1.0 | [0.9, 1] | 11.7 | [9.8, 13.5] |
| grazing rate | NA | poly2 | 15.4 | [8.4, 18.7] | 0.1 | [0, 0.1] | 9.5 | [7.1, 15.4] |

**Table S11)** Parameter estimates for TPC models fit to each isopod response. Both models included a quadratic (2nd-order polynomial) effect of temperature: $y = a + bT + cT^2$.

| **Response** | **Disease treatment** | **Model** | **a** | **95% CI** | **b** | **95% CI** | **c** | **95% CI** |
|---|---|---|---|---|---|---|---|---|
| isopod survival | diseased | binomial-poly2 | -6.7 | [-15.2, 3.6] | 1.3 | [0.1, 2.5] | 0.0 | [-0.1, 0] |
| isopod survival | healthy | binomial-poly2 | -30.4 | [-129.1, -5.8] | 4.8 | [1.2, 20.8] | -0.2 | [-0.7, 0] |
| grazing rate | NA | poly2 | -0.1 | [-0.2, 0.1] | 0.0 | [0, 0] | 0.0 | [0, 0] |

**Figure S10)** Hypothetical response of top-down interaction strength (effect of herbivores on net productivity) under scenarios with herbivores with contrasting thermal tolerance. Herbivore A is more sensitive to high temperatures than the producer and has the same thermal optimum ($T_{opt}$). Herbivore B is tolerant of high temperatures and has a $T_{opt}$ that is higher than the upper thermal limit ($T_{max}$) of the producer. **a)** Producer growth rate (productivity, *P*) vs. temperature in the absence of grazers. **b)** Grazing rates (*G*) vs. temperature of Herbivore A and B. **c)** Producer net growth rate (net productivity, $NP = P - G$) without grazers (green line) or with *either* Herbivore A (orange) or Herbivore B (purple). Eventually, the producer would go extinct at higher temperatures in the scenario with Herbivore B, as the herbivore consumes producer biomass faster than the producer can replace it. **d)** Top-down interaction strength ($\ln(NP_{alone}/NP_{+\ grazer})/t$) vs. temperature for scenarios with Herbivore A vs. with Herbivore B. Dashed lines indicate $T_{opt}$, and dotted lines indicate ($T_{max}$).

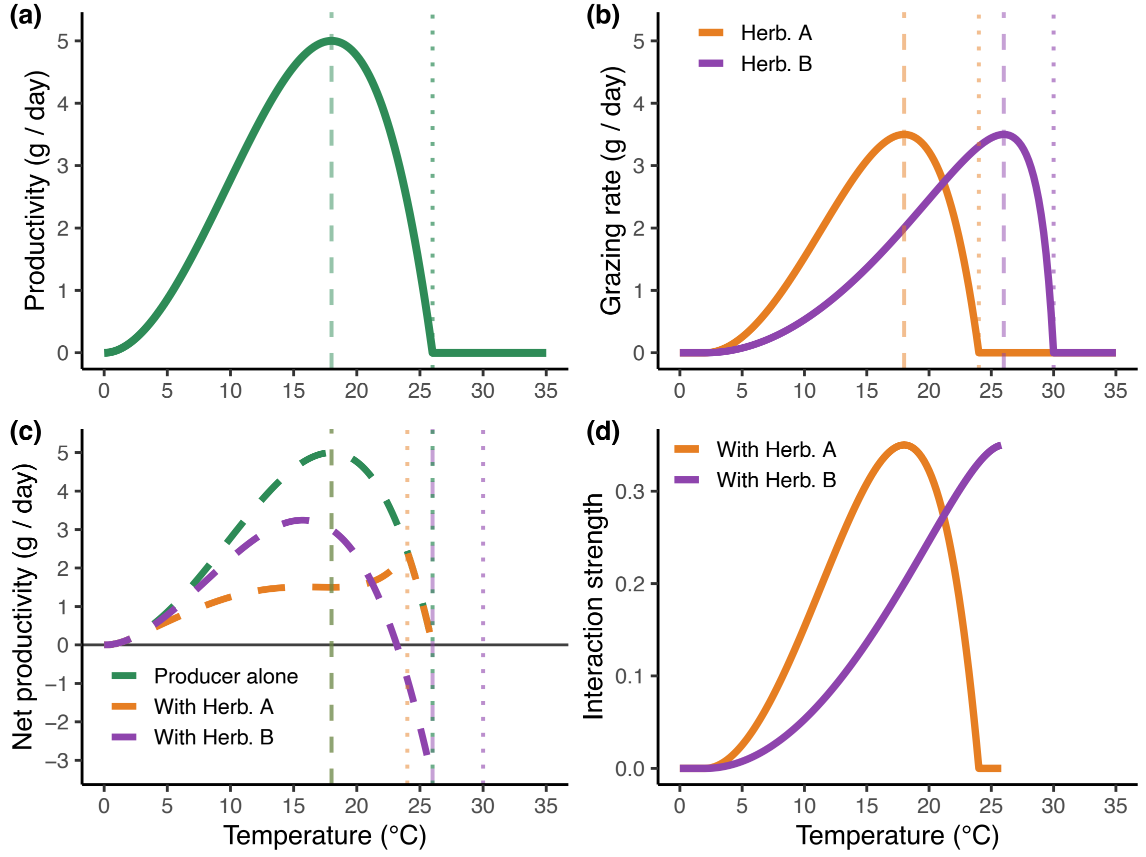
