## Supplemental model comparison tables for "Distinct thermal responses of a host plant and an invertebrate herbivore affect ecosystem productivity and disease dynamics in a coastal marine ecosystem"

for

**Description**: Model selection tables for different eelgrass and isopod response variables in the mesocosm and grazing experiments.

TABLE OF CONTENTS

Table S1: AICc model selection table for thermal performance curve (TPC) models fit to eelgrass responses in the no-grazing treatments. p. 2-3

Table S2: AICc model selection table for eelgrass responses in the grazing treatments. p. 4-5

Table S3: AICc model selection table for eelgrass disease onset time AFT models with different shape parameters. p. 6

Table S4: AICc model selection table for eelgrass disease onset time AFT models with different predictor structures. p. 6

Table S5: AICc model selection table for eelgrass final diseased area & disease severity. p 7

Table S6: AICc model selection table for GLM models fit to final diseased area. p. 8

Table S7: AICc model selection table for isopod TPC models. p. 9

**Table S1)** AICc model selection table for TPC models fit to eelgrass responses in the *no-grazing* treatments in the mesocosm experiment. The model used for predictions and parameter estimates is highlighted in **bold**.

| Response | Disease trt. | Grazing trt. | Model | AICc | ΔAICc | Rank |
|---|---|---|---|---|---|---|
| net growth | **diseased** | **no grazing** | **quadratic** | **-95.35** | **0** | **1** |
| | diseased | no grazing | lactin2 | -94.21 | 1.14 | 2 |
| | diseased | no grazing | weibull | -93.70 | 1.66 | 3 |
| | **healthy** | **no grazing** | **quadratic** | **-121.23** | **0** | **1** |
| | healthy | no grazing | lactin2 | -120.02 | 1.21 | 2 |
| | healthy | no grazing | weibull | -119.70 | 1.53 | 3 |
| leaf elongation | diseased | no grazing | quadratic | 74.64 | 0 | 1 |
| | **diseased** | **no grazing** | **weibull** | **75.69** | **1.05** | **2** |
| | diseased | no grazing | lactin2 | 76.16 | 1.52 | 3 |
| | **healthy** | **no grazing** | **weibull** | **72.80** | **0** | **1** |
| | healthy | no grazing | lactin2 | 75.33 | 2.53 | 2 |
| | healthy | no grazing | quadratic | 75.33 | 2.53 | 3 |
| leaf production | **diseased** | **no grazing** | **quadratic** | **-32.29** | **0** | **1** |
| | diseased | no grazing | lactin2 | -30.02 | 2.27 | 2 |
| | diseased | no grazing | weibull | -29.82 | 2.48 | 3 |
| | **healthy** | **no grazing** | **quadratic** | **-28.05** | **0** | **1** |
| | healthy | no grazing | weibull | -26.35 | 1.7 | 2 |
| | healthy | no grazing | lactin2 | -24.63 | 3.41 | 3 |
| rhizome dry mass | **diseased** | **no grazing** | **quadratic** | **-91.77** | **0** | **1** |
| | diseased | no grazing | weibull | -91.12 | 0.65 | 2 |
| | diseased | no grazing | lactin2 | -88.13 | 3.64 | 3 |
| | **healthy** | **no grazing** | **quadratic** | **-106.56** | **0** | **1** |
| | healthy | no grazing | weibull | -104.76 | 1.8 | 2 |
| | healthy | no grazing | lactin2 | -102.83 | 3.74 | 3 |
| aboveground dry mass | **diseased** | **no grazing** | **quadratic** | **-28.57** | **0** | **1** |
| | diseased | no grazing | weibull | -26.75 | 1.82 | 2 |
| | diseased | no grazing | lactin2 | -24.83 | 3.74 | 3 |
| | **healthy** | **no grazing** | **quadratic** | **-44.09** | **0** | **1** |
| | healthy | no grazing | weibull | -41.28 | 2.81 | 2 |
| | healthy | no grazing | lactin2 | -40.88 | 3.21 | 3 |
| belowground dry mass | **diseased** | **no grazing** | **quadratic** | **-74.27** | **0** | **1** |
| | diseased | no grazing | weibull | -71.26 | 3.01 | 2 |
| | diseased | no grazing | lactin2 | -71.02 | 3.25 | 3 |
| | **healthy** | **no grazing** | **weibull** | **-96.74** | **0** | **1** |
| | healthy | no grazing | quadratic | -96.46 | 0.28 | 2 |

| Response | Disease trt. | Grazing trt. | Model | AICc | ΔAICc | Rank |
|---|---|---|---|---|---|---|
| | healthy | no grazing | lactin2 | -89.81 | 6.94 | 3 |
| belowground: aboveground | **diseased** | **no grazing** | **quadratic** | **-39.80** | **0** | **1** |
| | diseased | no grazing | lactin2 | -38.14 | 1.66 | 2 |
| | diseased | no grazing | weibull | -37.38 | 2.42 | 3 |
| | **healthy** | **no grazing** | **quadratic** | **-32.18** | **0** | **1** |
| | healthy | no grazing | weibull | -31.28 | 0.9 | 2 |
| | healthy | no grazing | lactin2 | -27.70 | 4.48 | 3 |
| rhizome necroses | diseased | grazing | binomial-poly2 | 19.22 | - | - |
| | diseased | no grazing | binomial-poly2 | 23.96 | - | - |
| | healthy | grazing | binomial-poly2 | 21.76 | - | - |
| | healthy | no grazing | binomial-poly2 | 24.40 | - | - |

**Table S2)** AICc model selection table for non-linear models fit to eelgrass responses in the *grazing* treatments in the mesocosm experiment. The model used for predictions and parameter estimates is highlighted in **bold**.

| Eelgrass Response | Disease trt. | Grazing trt. | Model | AICc | ΔAICc | Rank |
|---|---|---|---|---|---|---|
| **net growth** | diseased | grazing | linear | -110.57 | 0 | 1 |
| | **diseased** | **grazing** | **log** | **-110.35** | **0.22** | **2** |
| | diseased | grazing | power | -107.88 | 2.69 | 3 |
| | diseased | grazing | asympReg | -107.87 | 2.7 | 4 |
| | **healthy** | **grazing** | **log** | **-74.19** | **0** | **1** |
| | healthy | grazing | linear | -74.13 | 0.05 | 2 |
| | healthy | grazing | asympReg | -71.54 | 2.65 | 3 |
| | healthy | grazing | power | -71.48 | 2.71 | 4 |
| **leaf elongation** | diseased | grazing | asympReg | 85.66 | 0 | 1 |
| | **diseased** | **grazing** | **log** | **85.95** | **0.29** | **2** |
| | diseased | grazing | linear | 87.48 | 1.82 | 3 |
| | diseased | grazing | power | 88.72 | 3.05 | 4 |
| | **healthy** | **grazing** | **log** | **86.41** | **0** | **1** |
| | healthy | grazing | linear | 86.80 | 0.39 | 2 |
| | healthy | grazing | asympReg | 89.08 | 2.67 | 3 |
| | healthy | grazing | power | 89.10 | 2.69 | 4 |
| **leaf production** | **diseased** | **grazing** | **log** | **-25.32** | **0** | **1** |
| | diseased | grazing | linear | -24.75 | 0.58 | 2 |
| | diseased | grazing | asympReg | -23.11 | 2.21 | 3 |
| | diseased | grazing | power | -22.59 | 2.73 | 4 |
| | **healthy** | **grazing** | **log** | **-28.35** | **0** | **1** |
| | healthy | grazing | linear | -28.10 | 0.25 | 2 |
| | healthy | grazing | asympReg | -25.71 | 2.63 | 3 |
| | healthy | grazing | power | -25.66 | 2.69 | 4 |
| **rhizome dry mass** | **diseased** | **grazing** | **log** | **-100.69** | **0** | **1** |
| | diseased | grazing | linear | -100.66 | 0.03 | 2 |
| | diseased | grazing | asympReg | -98.04 | 2.65 | 3 |
| | diseased | grazing | power | -97.98 | 2.71 | 4 |
| | **healthy** | **grazing** | **log** | **-104.29** | **0** | **1** |
| | healthy | grazing | linear | -103.99 | 0.3 | 2 |
| | healthy | grazing | asympReg | -102.29 | 2 | 3 |
| | healthy | grazing | power | -101.55 | 2.74 | 4 |
| **aboveground dry mass** | **diseased** | **grazing** | **log** | **-39.14** | **0** | **1** |
| | diseased | grazing | linear | -38.85 | 0.3 | 2 |
| | diseased | grazing | asympReg | -36.56 | 2.58 | 3 |

| Eelgrass Response | Disease trt. | Grazing trt. | Model | AICc | ΔAICc | Rank |
|---|---|---|---|---|---|---|
| | diseased | grazing | power | -36.42 | 2.72 | 4 |
| | **healthy** | **grazing** | **log** | **-43.26** | **0** | **1** |
| | healthy | grazing | linear | -42.54 | 0.71 | 2 |
| | healthy | grazing | asympReg | -41.54 | 1.72 | 3 |
| | healthy | grazing | power | -40.49 | 2.76 | 4 |
| **belowground dry mass** | diseased | grazing | linear | -88.30 | 0 | 1 |
| | **diseased** | **grazing** | **log** | **-88.00** | **0.3** | **2** |
| | diseased | grazing | power | -85.70 | 2.6 | 3 |
| | diseased | grazing | asympReg | -85.59 | 2.71 | 4 |
| | **healthy** | **grazing** | **log** | **-90.48** | **0** | **1** |
| | healthy | grazing | linear | -90.05 | 0.43 | 2 |
| | healthy | grazing | asympReg | -88.15 | 2.33 | 3 |
| | healthy | grazing | power | -87.73 | 2.76 | 4 |
| **belowground: aboveground** | diseased | grazing | linear | -40.59 | 0 | 1 |
| | **diseased** | **grazing** | **log** | **-40.43** | **0.16** | **2** |
| | diseased | grazing | power | -39.00 | 1.59 | 3 |
| | diseased | grazing | asympReg | -37.88 | 2.71 | 4 |
| | **healthy** | **grazing** | **log** | **-38.43** | **0** | **1** |
| | healthy | grazing | linear | -38.40 | 0.04 | 2 |
| | healthy | grazing | asympReg | -35.81 | 2.63 | 3 |
| | healthy | grazing | power | -35.80 | 2.63 | 4 |

**Table S3** AICc model selection table for eelgrass disease onset time, comparing accelerated failure time (AFT) models with different shape parameters (Weibull, log-normal, log-logistic). Models all had the same additive predictor structure (no interactions between the predictors: temperature, disease treatment, grazing treatment). The best model is highlighted in **bold**.

| Response | Model | k | AICc | ΔAICc | Rank |
|---|---|---|---|---|---|
| disease onset | **additive: Weibull** | **5** | **605.99** | **0.00** | **1** |
| | additive: log-normal | 5 | 613.03 | 7.05 | 2 |
| | additive: log-logistic | 5 | 617.14 | 11.16 | 3 |

**Table S4** AICc model selection table for eelgrass disease onset time, comparing accelerated failure time (AFT) models with the same shape parameter (Weibull), but varied predictor structure (temperature, disease treatment, grazing treatment). The best model is highlighted in **bold**.

| Response | Model | k | AICc | ΔAICc | Rank |
|---|---|---|---|---|---|
| disease onset | **disease.trt + grazing.trt + temp** | **5** | **605.99** | **0.00** | **1** |
| | disease.trt x grazing.trt + temp | 6 | 607.34 | 1.36 | 2 |
| | disease.trt + grazing.trt x temp | 6 | 607.97 | 1.99 | 3 |
| | disease.trt x temp + grazing.trt | 6 | 608.21 | 2.23 | 4 |
| | disease.trt x temp x grazing.trt | 9 | 613.90 | 7.92 | 5 |

**Table S5** AICc model selection table for non-linear models fit to eelgrass final diseased area and disease severity data. The model that was used for predictions and parameter estimates are highlighted in **bold.**

| Response | Disease trt. | Grazing trt. | Model | AICc | ΔAICc | Rank |
|---|---|---|---|---|---|---|
| diseased area | **diseased** | **grazing** | **exponential** | **376.28** | **0** | **1** |
| | diseased | grazing | linear | 376.83 | 0.55 | 2 |
| | diseased | grazing | polynomial2 | 378.90 | 2.62 | 3 |
| | diseased | grazing | power | 378.94 | 2.66 | 4 |
| | diseased | no grazing | linear | 334.48 | 0 | 1 |
| | **diseased** | **no grazing** | **exponential** | **334.88** | **0.4** | **2** |
| | diseased | no grazing | polynomial2 | 337.08 | 2.6 | 3 |
| | diseased | no grazing | power | 337.14 | 2.66 | 4 |
| | **healthy** | **grazing** | **exponential** | **392.26** | **0** | **1** |
| | healthy | grazing | power | 393.58 | 1.33 | 2 |
| | healthy | grazing | polynomial2 | 395.73 | 3.47 | 3 |
| | healthy | grazing | linear | 395.91 | 3.66 | 4 |
| | **healthy** | **no grazing** | **exponential** | **397.18** | **0** | **1** |
| | healthy | no grazing | linear | 397.75 | 0.57 | 2 |
| | healthy | no grazing | polynomial2 | 399.37 | 2.18 | 3 |
| | healthy | no grazing | power | 399.57 | 2.39 | 4 |
| disease severity | **diseased** | **grazing** | **exponential** | **-123.90** | **0** | **1** |
| | diseased | grazing | linear | -123.77 | 0.12 | 2 |
| | diseased | grazing | polynomial2 | -121.33 | 2.56 | 3 |
| | diseased | grazing | power | -121.30 | 2.59 | 4 |
| | **diseased** | **no grazing** | **linear** | **-189.29** | **0** | **1** |
| | diseased | no grazing | exponential | -189.15 | 0.15 | 2 |
| | diseased | no grazing | power | -186.64 | 2.65 | 3 |
| | diseased | no grazing | polynomial2 | -186.64 | 2.65 | 4 |
| | **healthy** | **grazing** | **exponential** | **-107.11** | **0** | **1** |
| | healthy | grazing | linear | -106.83 | 0.28 | 2 |
| | healthy | grazing | polynomial2 | -106.80 | 0.31 | 3 |
| | healthy | grazing | power | -106.34 | 0.77 | 4 |
| | **healthy** | **no grazing** | **exponential** | **-126.04** | **0** | **1** |
| | healthy | no grazing | linear | -125.97 | 0.06 | 2 |
| | healthy | no grazing | polynomial2 | -125.37 | 0.67 | 3 |
| | healthy | no grazing | power | -124.16 | 1.88 | 4 |

**Table S6)** AICc model selection table for Gamma GLM models (log link) fit to eelgrass final diseased area. The best-supported model is highlighted in **bold**.

| Response | Model | k | AICc | ΔAICc | Rank |
|---|---|---|---|---|---|
| diseased area | **disease x grazing + temp** | **6** | **1,432.11** | **0.00** | **1** |
| | disease + grazing + temp | 5 | 1,432.39 | 0.29 | 2 |
| | grazing x temp + disease | 6 | 1,433.75 | 1.65 | 3 |
| | disease x temp + grazing | 6 | 1,434.61 | 2.50 | 4 |
| | disease x grazing x temp | 9 | 1,437.78 | 5.67 | 5 |

**Table S7)** Model comparison table for different TPC models fit to each isopod response. The model that was used for predictions and parameter estimates are highlighted in **bold.**

| Response | Disease trt. | Model | AICc | ΔAICc | Rank |
|---|---|---|---|---|---|
| isopod survival | healthy | **binomial-poly2** | **49.09** | - | - |
| | diseased | **binomial-poly2** | **57.29** | - | - |
| grazing rate | NA | **poly2** | **-250.23** | **0** | **1** |
| | NA | lactin2 | -250.17 | 0.06 | 2 |
| | NA | weibull | -249.18 | 1.06 | 3 |
